# Heterogeneity and variability in short term synaptic plasticity and implications for signal transformation

**DOI:** 10.1101/2025.06.28.662098

**Authors:** Sulu Mohan, Upinder Singh Bhalla

## Abstract

There are many sources of heterogeneity in the CA1 network, including plasticity, connectivity and cell properties, yet the extent and functional consequences of this diversity remains poorly understood. We used patterned optogenetic stimulation of CA3 pyramidal neurons and whole-cell patch clamp recordings from CA1 pyramidal neurons in acute mouse hippocampal slices, to characterize the contributions of different forms of heterogeneity to information flow. We found pronounced heterogeneity in synaptic responses and short-term plasticity (STP), influenced by the neurotransmitter identity, input pattern, and size of the activated presynaptic ensemble. Inhibitory synapses exhibited greater diversity in both response variability and depression profiles than excitatory synapses. We incorporated these readings in a molecule-to-network multiscale model of the CA3->CA1 circuit. The reference model shows strong decorrelation of autocorrelated input, but removal of STP makes the decorrelation frequency dependent. Removal of stochasticity and heterogeneity in connections makes the output periodic. Thus heterogeneity, short-term plasticity, and stochasticity each have distinct effects on cellular information transmission.

## Introduction

Short term synaptic plasticity (STP) is the activity dependent modulation of synaptic strengths in the time scale of milliseconds to seconds (Abbott & Regehr, 2004; Regehr, 2012; Thomson, 2000; Zucker & Regehr, 2002a). STP in the hippocampus is crucial for encoding memory, guiding spatial navigation, and processing temporal patterns of inputs (Deng & Klyachko, 2011; Zucker & Regehr, 2002b). STP is a diverse phenomenon with distinct profiles not only across brain regions and cell types but even among synapses formed by the same presynaptic neuron onto distinct postsynaptic targets (Larsen & Sjöström, 2015; Markram et al., 1998; Reyes et al., 1998). This raises the question of whether such diversity is a bug - to be worked around, or a feature - providing functional benefits.

Synaptic diversity has been documented across cortical and hippocampal circuits. Excitatory projections of the same CA3 neuron, for instance, show different STP based on their postsynaptic targets - short-term depressing when synapsing onto a subset of interneurons while facilitating when synapsing onto CA1 pyramidal cells (Sun et al., 2018a). Likewise, studies in thalamocortical and motor cortex circuits have shown cell-to-cell variability in STP within the same cell type (Díaz-Quesada et al., 2014a; McFarlan et al., 2024). Even synapses within the same neuron have been shown to have heterogeneous STP in hippocampal CA1 (Dobrunz & Stevens, 1997a) and mouse barrel cortex (Buchholz et al., 2023a) neurons. Dobrunz & Stevens, 1997, demonstrated that synaptic heterogeneity in STP can be explained by the wide release probability range (0.05 – 0.9 for the first pulse, 0.2 – 0.9 for the second pulse) even among individual CA1 neuron synapses. This low average probability of release (0.2) also makes CA1 synapses highly stochastic (Branco & Staras, 2009; Murthy et al., 1997). Our study extends these findings in physiologically relevant network conditions involving both excitation and inhibition. Previous studies using minimal stimulation or single-pathway activation do not incorporate complex network dynamics, such as feedforward inhibition or ensemble input integration.

Heterogeneity and variability are not merely sources of noise, but computationally meaningful features with implications that extend from synaptic to behavioural levels. Computational models rely on synaptic heterogeneity to reproduce *in vivo*-like spike timing and network dynamics (Buchholz et al., 2023a; Hass et al., 2016). Heterogeneous networks show improved learning and information processing, especially for tasks with a rich temporal structure (Di Volo & Destexhe, 2021; Perez-Nieves et al., 2021; Rotman & Klyachko, 2013). In attractor network models of working memory, incorporating realistic STP diversity improves the biological accuracy of hippocampal and prefrontal simulations, supports multiple memory states, and improves sensitivity to input timing (Barak & Tsodyks, 2007; Hass et al., 2022). Behavioral experiments in awake electric fish have linked differences in short-term depression to directional tuning during movement (Chacron et al., 2009). These findings underscore the importance of synaptic variability in enabling flexible and robust computation in neural circuits.

In the hippocampus, this heterogeneity becomes especially important in the transformation of CA3 inputs by CA1 neurons. It has been shown that short-term depression of stochastic CA1 synapses reduces input redundancy by decorrelating spike trains, making postsynaptic responses less predictable and more efficient (Abbott et al., 1997). Short-term facilitation, on the other hand, amplifies high-frequency bursting inputs that are essential for CA1 functions such as place cell formation (Izhikevich et al., 2003; Lisman, 1997). The stochasticity resulting from low release probability further introduces trial-to-trial variability, allowing CA1 to become selectively sensitive to both temporal and rate-based features of the input (Rodrigues et al., 2023). These studies show that CA1 STP actively transforms inputs using plasticity and probabilistic transmission and does not just passively filter, gate and integrate them. Our work uses spatially patterned input to add a further dimension to earlier studies and to systematically delineate effects of stochasticity and STP on input-output transformations in CA1. Using a detailed computational model of the CA3-CA1 network that can take patterned inputs of varying stimulus strength and frequencies, we investigated the frequency-dependent transformation of autocorrelated inputs with and without stochasticity, STP and inhibition.

## Results

Our study was structured from subcellular to network scales. At each level we checked computational model performance against experiments, as follows: First we characterized heterogeneity in synaptic transmission and STP at the cell level. We then examined response diversity to combinations of inputs. Next we estimated synapse-level heterogeneity and stochasticity. We then stepped back to look at cellular summation. Finally we used the model to examine decorrelation of input spiking patterns from CA3 to CA1, and dissect contributions from connection heterogeneity, STP, and stochasticity to this input-output transform.

### Variability in CA1 responses is not explained by stimulus variability

We analyzed the overall variability in CA1 responses and short term plasticity at two levels: 1) cell and synapse heterogeneity, which refers to intrinsic differences in cellular and synaptic properties, and 2) trial-to-trial variability over multiple repeats of the same stimulus (Figure 1A). We used Grik-Cre mice with CA3 specific expression of channelrhodopsin (ChR2) upon injection of lox-ChR2 virus. We used a Digital Mirror Device (Polygon 400) to deliver patterned optical stimuli (470 nm wavelength) onto the CA3 region of acute transverse hippocampal slices with intact CA3–CA1 feedforward excitation-inhibition circuitry, while CA1 responses were recorded using whole cell patch clamp (Figure 1B, C). Each individual square (spot size 13 x 7 µm, power 14.5 µW/spot) in the optical patterns is expected to reliably elicit a single spike at the target CA3 cell (Bhatia et al., 2019). A field recording electrode placed close to the CA3 cell body layer acted as a control for measuring variability at the level of stimulation (Figure 1B). Optical patterns consisted of 1, 5 or 15 simultaneously active randomly arranged non-overlapping squares. We measured the excitatory component of the postsynaptic response by holding the CA1 neuron at -70 mV, the reversal for inhibition, and in the same experiment measured inhibitory postsynaptic currents by holding the cell at 0 mV, the reversal for excitation. STP was measured by pairing the same optical pattern at 50 ms interval (Figure 1D). STP ratio (STPR) was calculated as the difference in peak amplitudes normalised to the sum of peaks. In order to test the effects of summation of inputs, optical patterns were designed such that each 5-square pattern is a composite of five 1-square patterns and the 15-square pattern is a composite of three 5-square patterns (Figure 1E).

**Figure 1:**
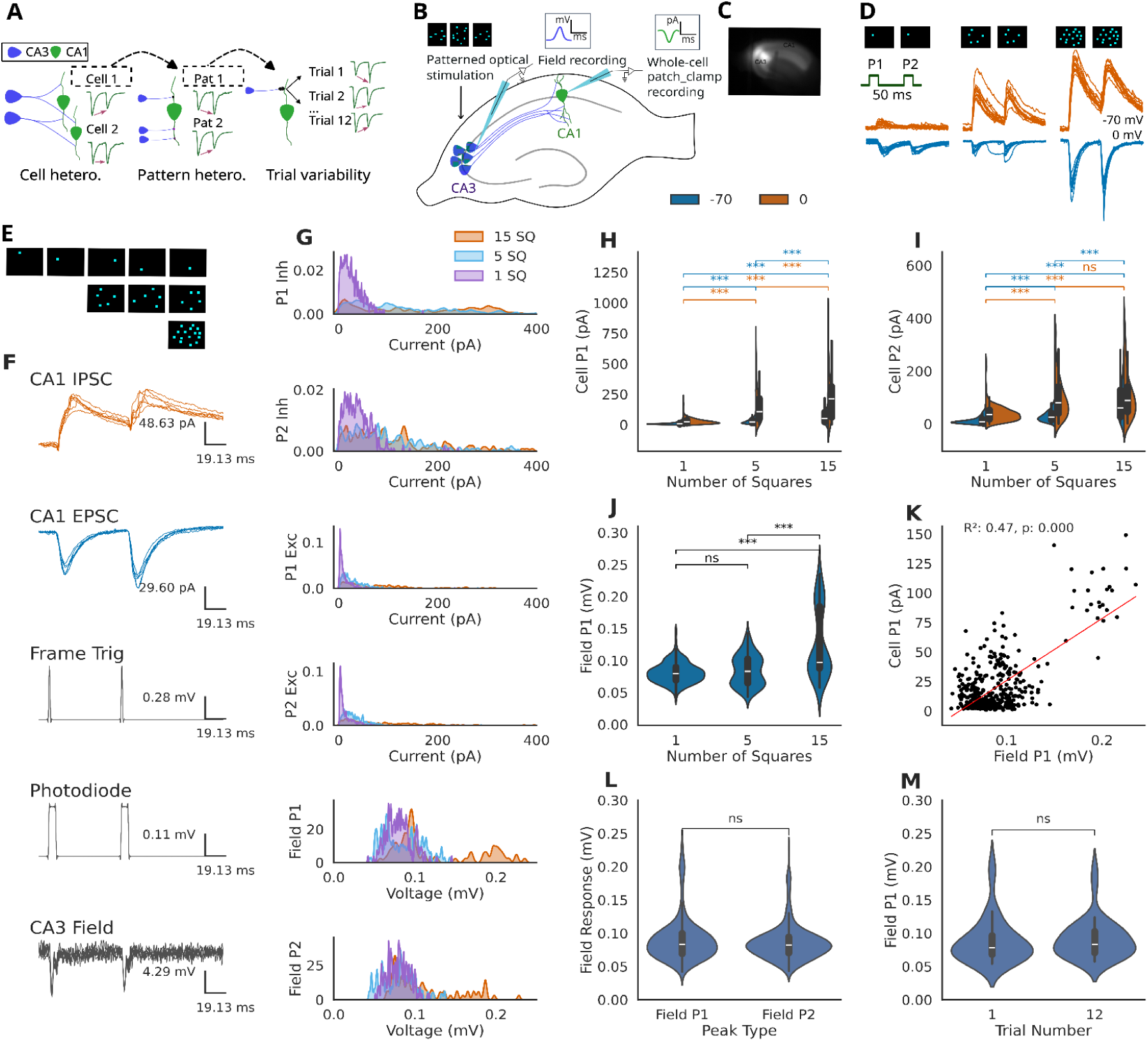
Variability in CA1 responses is not explained by stimulus variability. A. Schematic illustrating cellular heterogeneity, referring to between-cell variability; pattern heterogeneity, referring to within-cell variability reflected in responses to distinct optical patterns; and trial-to-trial variability, referring to fluctuations in response when repeating the same pattern (12 repeats). B. Schematic showing acute transverse slices of mouse hippocampus expressing channelrhodopsin specifically in CA3. Optogenetic activation of CA3 was delivered using a DMD device (Polygon) integrated into the light path of the microscope. CA3 activity was recorded with a field electrode placed near the cell body layer, while CA1 responses were recorded using whole-cell patch clamp. C. Fluorescence image showing channelrhodopsin expression (Td-Tomato) in CA3. CA3 provides direct excitatory input and feedforward inhibition to CA1. D. Optical patterns were either 1, 5 or 15 non-overlapping squares, randomly chosen in a 24x24 optical grid. STP was tested by pairing the same optical pattern at 50 ms interval. CA1 was voltage clamped at -70 mV for recording Excitatory PostSynaptic Currents (EPSC traces in blue) and at 0 mV for recording Inhibitory PostSynaptic Currents (IPSC traces in red). E. There were fifteen 1-square patterns which were then clubbed into three 5-square and one 15-square pattern. F. Representative raw traces from a single CA1 pyramidal cell recording in response to a 15 square (15 SQ) stimulation pattern. From top to bottom: CA1 inhibitory postsynaptic current (IPSC, orange) recorded at 0 mV; CA1 excitatory postsynaptic current (EPSC, blue) recorded at -70 mV; frame trigger to switch optical patterns; photodiode signal for stimulus timing; and the corresponding local field potential recorded in the CA3 pyramidal layer. G. Ridgeline density plots showing the distribution of response amplitudes across the entire population for the first (P1) and second (P2) peaks. Distributions are shown for inhibitory currents (P1 and P2 Inh), excitatory currents (P1 and P2 Exc), and CA3 field potentials (Field P1 and P2). Responses to 1, 5, and 15 SQ stimuli are color-coded as indicated in the legend. The probability density was estimated using a Gaussian Kernel Density Estimation (KDE) with a bandwidth adjustment factor of 0.1. H. Summary of peak synaptic currents across all recorded cells. Split violin plots show the distribution of the first peak amplitudes as a function of the number of stimulated squares (1, 5, 15 Squares). Orange and blue represent IPSCs and EPSCs, respectively. Statistical comparisons between 1, 5 and 15 patterns are shown (Mann-Whitney U-test: *p<0.05, **p<0.01, ***p<0.001). I. Same as H for second peak amplitudes J. Summary of CA3 field potential amplitude (1, 5, 15 SQ). The violin plot shows the distribution of the first peak-to-trough amplitude (Field P1) for selected cells. K. Correlation between EPSCs and local field potentials. The scatter plot shows the relationship between Cell P1 and Field P1. The red line indicates the linear regression, with the coefficient of determination (R^2^) and p-value shown. L. Desensitization of CA3 field potential: distribution of the first peak (Field P1) and second peak (Field P2) amplitudes across all conditions M. Stability of the field potential response over time: Field potential peak (Field P1) between the first and last trials (Trial 1 vs. Trial 12) of the experiment.

To control for variability at the level of stimulation, we recorded field responses using an electrode placed in CA3. Field responses were biphasic for most experiments and therefore peak-to-peak amplitudes were measured. Field responses did not show significant differences between paired pulses (Figure 1L) and across different trials (Figure 1M). This allowed us to rule out ChR2 desensitization as a potential source of observed variability in synaptic responses.

### Cells are heterogenous in responses and short term plasticity

To test for heterogeneity between cells in synaptic and STP properties, we performed voltage clamp recordings in CA1 cells (n=18). We first checked that global physiological properties of holding current (Figure 2A) and input resistance (Figure 2B) were within physiological ranges and stable over the experiment. We applied additional criteria for signal-to-noise for segments of the recording at highly depolarized holding potentials. We then also computed the IE ratio. This was consistent across cells (Figure 2C).

**Figure 2:**
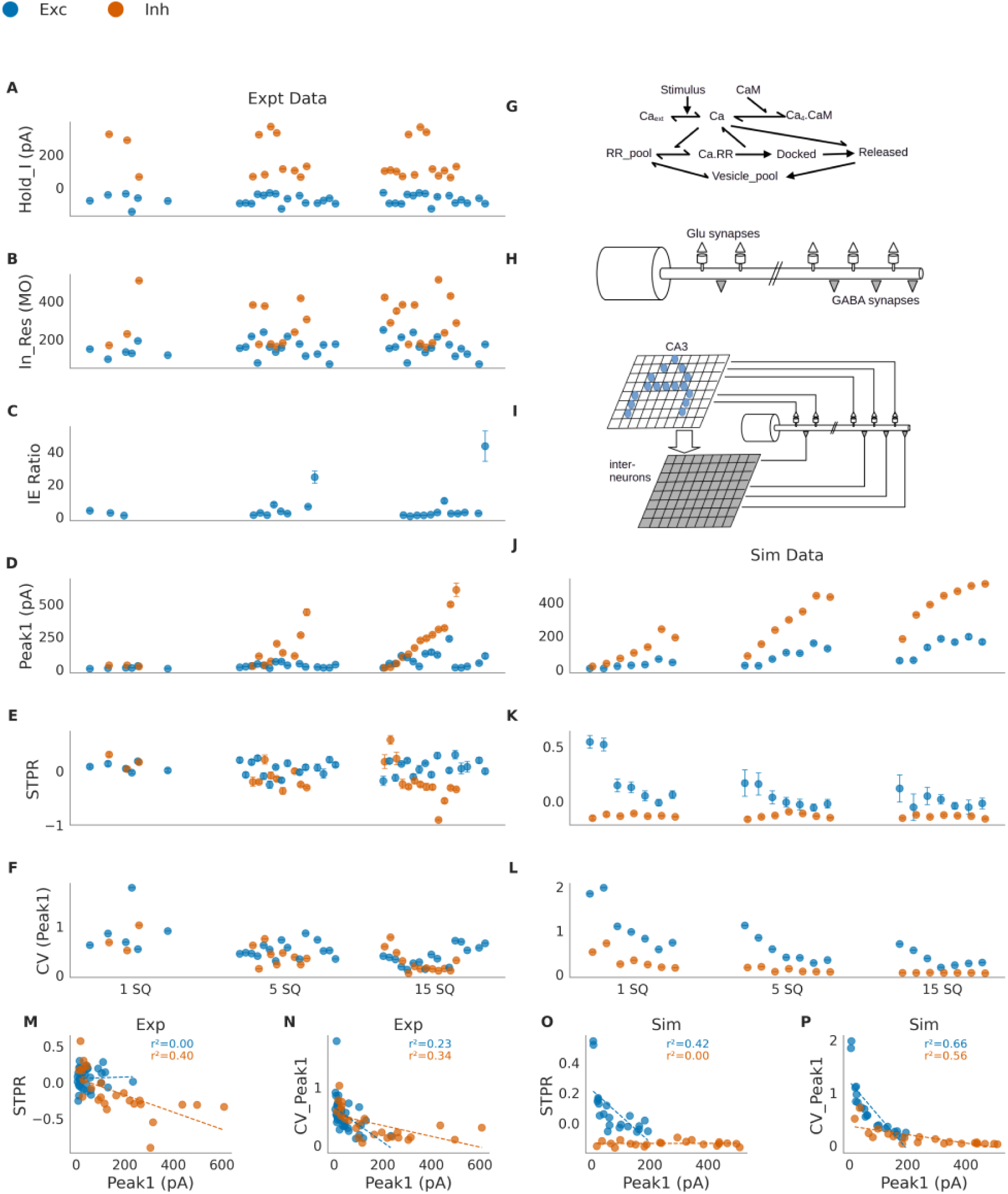
Cells are heterogenous in responses and short term plasticity. Cells with data on both excitation and inhibition are sorted in the ascending order of first peak amplitude of IPSCs for the 15 square pattern (see panel D). Blue dots represent excitation, orange dots represent inhibition. Responses are grouped by 1, 5, and 15 square patterns along the x-axis. In row order: A. Mean holding current for cells (pA) B. Mean input resistance for recorded cells (MOhm) C. Ratio of inhibitory to excitatory first peak amplitudes D. Mean first peak amplitude (pA) E. Short term plasticity ratio (STPR) calculated as (peak2-peak1)/(peak2+peak1) F. Coefficient of variation for peak1 (G, H, I) Model architecture. The model was based closely on (Asopa & Bhalla, 2025). G. Reaction scheme for presynaptic neurotransmitter release. The same scheme was used for glutamatergic and GABAergic synapses, but the rates differed. H. Schematic of spiking neuronal model. The cell was modeled as a 10-micron soma with a 200 micron long, 2 micron diameter dendrite. Glu synapses terminated on 0.5 micron diameter dendritic spines at a mean spacing of 2 microns. GABAergic synapses terminated on the dendrite with a mean spacing of 1 micron. Each presynaptic bouton for Glu as well as GABA had an independent reaction scheme for neurotransmitter release. I. Schematic of input and network. A 16x16 array of single-compartment CA3 neurons received patterned optical stimuli. They projected onto a 16x16 array of binary threshold interneurons using a random connectivity matrix (RCM). The CA3 neurons projected onto CA1 Glu synapses using a random connectivity matrix, and as binary excitatory input onto the interneurons using a distinct RCM. The interneurons in turn projected onto the GABAergic synapses of the CA1 neuron using another RCM. J. Simulation corresponding to panel D. The trends observed in experimental peak 1 data were reproduced by varying the pattern density parameter. Pattern density refers to the number of simulated CA cells for any given spot of stimulation. These ‘cells’ are sorted on the basis of ascending order of their peak 1 inhibitory amplitudes. K. STPR sorted in the same order as J for simulated data. L. CV sorted in the same order as J for simulated data. Note decline in CV with increasing number pattern density and number of input squares. M. Linear regression plots showing negative correlation of STPR with PSC amplitudes only for inhibition (R^2^ = 0.40, p<0.001). N. Linear regression plots showing negative correlation of CV with number of squares in pattern and PSC amplitudes for both excitation (R^2^=0.23, p=0.002) and inhibition (R^2^=0.34, p=0.002). O. Simulated data for pattern density distribution does not show the negative correlation of STPR with weights. P. Simulated data for pattern density distribution accurately replicated the decrease in CV with increasing number of spots as well as peak 1 amplitudes for a given stimulus strength.

Next we investigated the heterogeneity of E/IPSCs (Figure 2D) between cells. For clarity of comparison, all graphs from A to F use the cell-ordering obtained from sorting IPSC amplitudes measured for the 15-square patterns (Figure 2D). There was a wide range of IPSC and EPSC amplitudes and as expected from Figure 2C the correlation was only modest. We defined the short-term plasticity ratio as STPR = (peak2-peak1)/(peak2+peak1). STPR for excitatory (E) synapses was scattered in a narrow band near zero, but for inhibitory (I) synapses there was a wider range centered below zero (Figure 2E). As expected, the trial to trial variability (coefficient of variation in multiple repeats of the same pattern) declined with the number of illuminated squares (Figure 2F). This heterogeneity in facilitation of excitatory and depression of inhibitory inputs to CA1 neurons is generally masked when STP is measured by volley electrical stimulation of Schaffer collaterals (Supplementary Figure 1), which is how it is conventionally measured (Dobrunz & Stevens, 1997a; Jackman & Regehr, 2017; Zucker & Regehr, 2002b).

We then used our data to fine-tune a previously developed model of the CA3-CA1 network designed for exactly the same optical stimulus and patch recording configuration (Asopa & Bhalla, 2025). In brief, the model included stochastic chemical signaling for both E and I presynaptic boutons, a detailed compartmental conductance-based CA1 neuron model for cellular electrophysiology, an array of interneurons, and an array of CA3 neurons coupled to the optical stimulus (Figure 2G-I, methods).

As a first tuning step, we tested how to obtain differential responses between cell recordings. While systematically changing synaptic conductances between models could trivially account for different amplitude responses, it was neither physiologically plausible nor was able to explain trial-to-trial variability (data not shown). We instead tested if we could account for the response distributions by simulating the known differences of ChR2 expression levels between slices. We did so by implementing a ‘pattern density’ parameter in our model, where pattern density refers to the number of CA3 cells stimulated by a given light spot. Thus, our panel of model cells was formed by choosing pattern density values between 1 and 16. With this set of model cells, we were able to replicate the range and relative values of the Peak 1 E and I responses (Figure 2J). Our model also replicated the relative STPR ratios of E and I synapses but not the scatter between observations (Figure 2K). Importantly, the decline in CV between trials was replicated by the model without any tuning (Figure 2L).

Numerous studies suggest a negative relationship between plasticity and synaptic strength (Dobrunz & Stevens, 1997a; Thomson, 2000; Zucker & Regehr, 2002b). We examined this in our dataset by plotting STPR against peak 1. While there was a weak (R2=0.40, p<0.001) correlation for the I synapses, E synapses showed no dependence (R2=0.00, p=0.80) (Figure 2M).

The CV between trials is an indirect estimate of the number of synapses activated per illuminated square, since larger numbers of synapses will average out and give a smaller CV over repeated trials. If this is the case, then recordings with stronger expression should have larger E(I)PSCs and smaller CV. We found that this was indeed the case, though the strength of the effect was small (R2=0.23, p=0.003 for Exc, R2=0.334, p=0.002 for Inh), (Figure 2N). The model, without tweaking, agreed on all these points except for a higher STPR for weak excitatory synapses and lower STPR for strong inhibitory synapses (Figure 2M, O).

Thus at this stage we had characterized synaptic current amplitudes and STPR across cells, and found considerable heterogeneity. We were able to replicate most of the heterogeneity in a model by assuming that expression levels, and hence ChR2 activated CA3 cell numbers differed. This interpretation was consistent with the observed reduction in CV with stronger E/IPSC responses.

### Cells show clustering on the basis of STP ratios

How similar are cells in their STP ratios? To check this, we performed bootstrap resampling (10000 iterations) from observed STPR data for all possible cell pair combinations and calculated the p-value of the difference between each cell pair. This was done for four different conditions: 5-square excitation, 15-square excitation, 5-square inhibition and 15-square inhibition, for both experimental and simulated datasets. We compared cell pairs for similarity in 5-square STP ratios and found a broad distribution with some cell pairs showing high similarity (p=0.95; Figure 3A, E, C, G) and others with marked dissimilarity (p=1.00e-5; Figure 3B, F, D, H). This was observed for both excitation and inhibition in experimental data (Figure 3A-D) and to a lesser extent in simulated data (Figure 3E-H).

**Figure 3:**
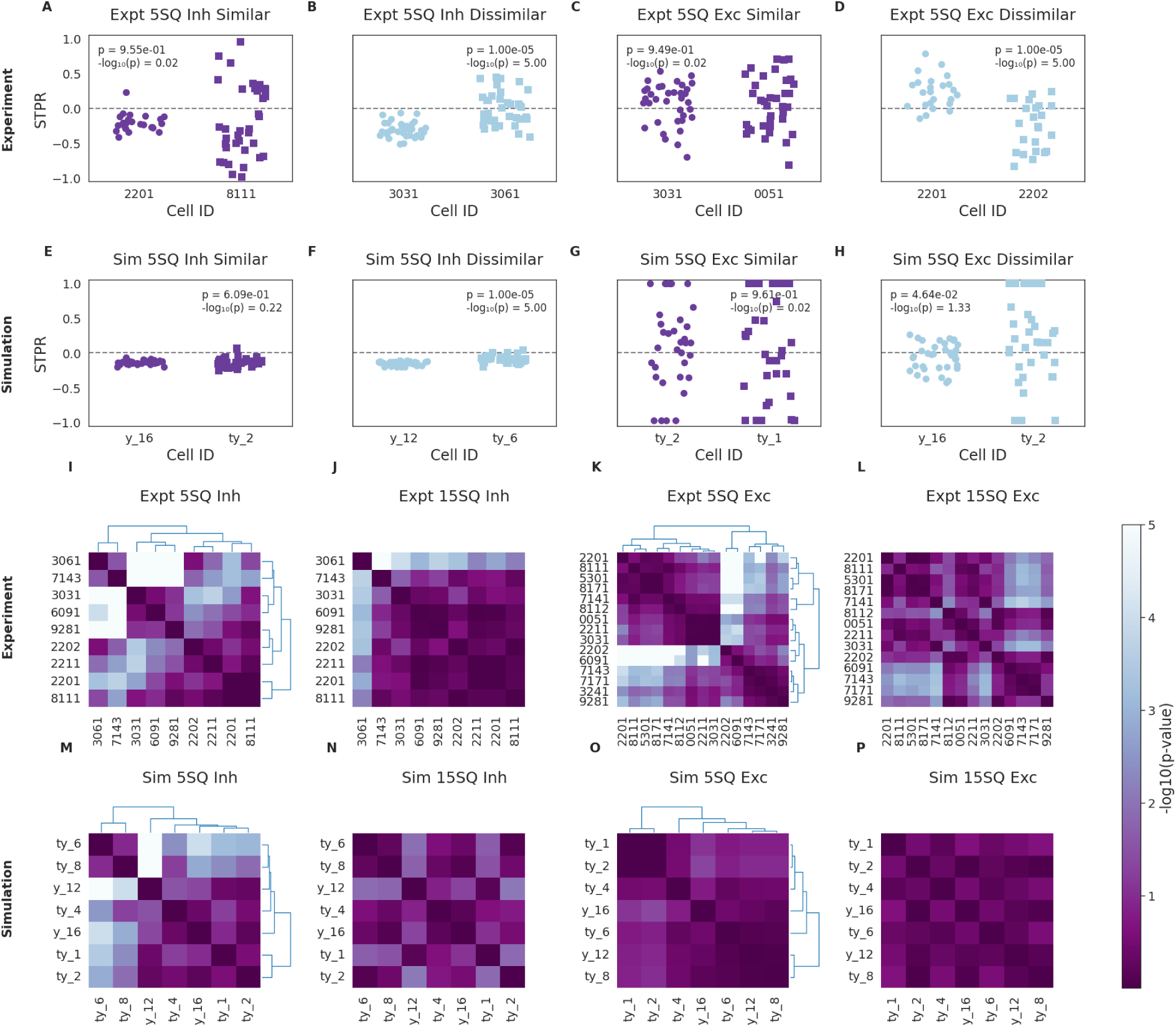
Cells form distinct clusters based on their STP profiles. Clustered heat maps showing clustering of cells based on the differences between their STP ratios across four conditions: 5-square inhibition, 15-square inhibition, 5-square excitation, and 15-square excitation. The color intensity in each cell of the heatmap represents the negative log of the p-value, which quantifies the significance of the mean STPR difference for a specific cell pair. This p-value is derived by comparing the actual observed mean STPR difference to a distribution of mean differences obtained through 10000 bootstrap resampling iterations from that cell pair. Essentially, it indicates the probability that the observed STPR difference occurred by chance. Dendrograms, displayed to the right and top of 5-square heatmaps represent hierarchical clustering based on Euclidean distance between the cells’ STPR profiles. Darker colors highlight cell pairs with more significant STPR differences (i.e., lower p-values). (A-H) STPR Distributions of Representative Cell Pairs. Stripplots of STPR values for the most similar and most dissimilar cell pairs, as determined for the 5-square patterns. Dots represent trials. The inset text shows the p-value and the corresponding -log10(p-value) for the comparison. Experimental data: Similar (A) and dissimilar (D) pairs for the 5SQ Inhibition. Similar (C) and dissimilar (D) pairs for the 5SQ Excitation. (E-H) Same as A-D but for simulation data (I-P) Heatmap for pairwise cell comparison based on STPR, heat represents the -log10(p-value). Darker colors indicate greater dissimilarity (i.e., a lower p-value, capped at p < 1e-5). Hierarchical clustering was performed on the 5SQ data (I, K, M, O) to group cells based on their STPR similarity and to generate the dendrograms shown. The resulting cell order is then used to plot corresponding 15SQ data (J, L, N, P). Experimental Data Heatmaps for 5SQ Inhibition, clustered (I), 15SQ Inhibition, ordered as in I (J), 5SQ Excitation, clustered (K), 15SQ Excitation, ordered as in K (L). Same as I-L but for simulation (M-P)

Do cells cluster on the basis of STPR in 5-square patterns? Hierarchical clustering was performed on the Euclidean distance between the -log10(p_values) obtained using the bootstrap approach. The resulting -log10 (p-value) matrices derived from 5-square data were used to establish a primary cell ordering. This 5-square derived order was subsequently applied to visualise heatmaps for both 5 and 15-square patterns, enabling comparison across pattern sizes. The dendrograms show the clustering for the 5-square pattern (Figure 3I, K). We found that both excitatory and inhibitory STPR responses did indeed form clusters which were mostly consistent between 5 and 15 square comparisons (Figure 3I-L). There were similar but weaker outcomes of clustering for the simulated datasets (Figure 3M-P).

Pairwise similarity was further assessed using two other methods: (1) using an ANOVA-based approach (Supplementary Figure 2) and (2) by computing the Euclidean distance between cell pairs in a 2-dimensional space defined by the mean and standard deviation of STPR (Supplementary Figure 3). This latter approach revealed that cells clustered tightly in this space for simulated cells in inhibition (Supplementary Figure 3Q, R) whereas excitation showed marked similarity in distribution for experimental and simulated data (Supplementary Figure 3S, T). Thus this analysis showed that there was some clustering of recorded cell STPR values both for 5 and 15 square patterns.

### Distinct spatial input patterns elicit distinct inhibitory responses

We next investigated heterogeneity at the level of spatial patterns of synaptic input. We asked whether distinct 5-square spatial patterns evoke heterogeneous E(I)PSC responses in CA1 neurons, and also heterogeneity in STP over the corresponding patterns. Three different 5-square patterns were tested per cell, with 12 repeats per pattern. We performed pairwise comparisons of E(I)PSCS between patterns within each cell, assessing significance using bootstrap resampling (1000 iterations). We found that IPSCs had greater heterogeneity between patterns than EPSCs, and this was also seen in the simulations without any tuning (Figure 4A-F). Similarly, STPR was more heterogeneous for IPSCs than for EPSCs (Figure 4G-K). These effects are summarized in Figure 4M-N.

**Figure 4:**
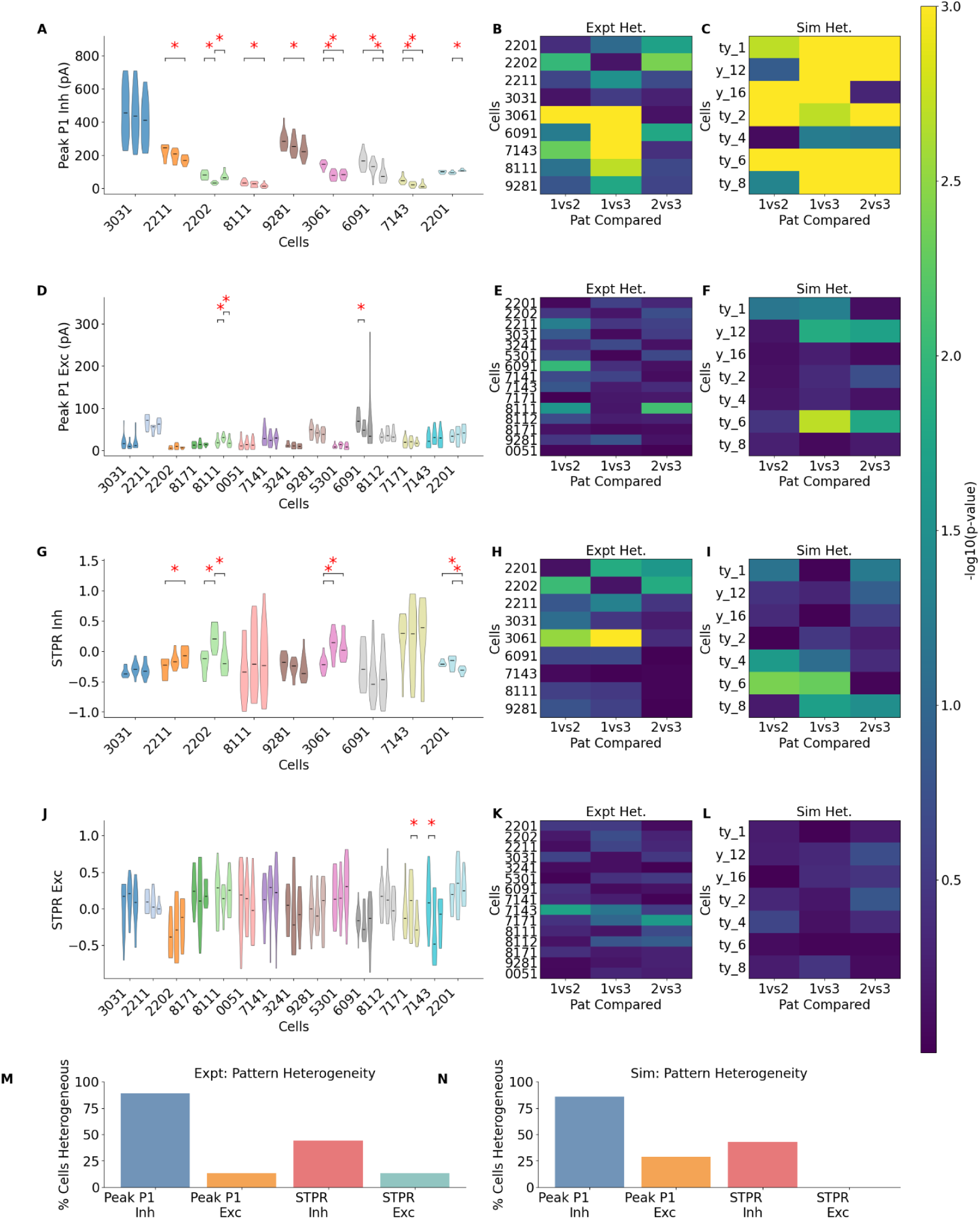
Pattern heterogeneity in initial responses are not reflected in their STP ratios. Heterogeneity of initial synaptic responses and STPR responses to 3 different 5-square patterns across experimental and simulated cells. (A-C) Peak 1 Amplitude during Inhibition. A. Experimental distributions of Peak 1 for three patterns per cell (each cell uniquely colored). Asterisks denote p < 0.05 for pattern-pair differences within a cell. B. Experimental heatmap: -log10 (p-value) for Peak 1 differences between pattern pairs (1v2, 1v3, 2v3) within each cell. C. Simulation heatmap, analogous to (B), for Peak 1 during inhibition. (D-F) Peak 1 Amplitude during Excitation D. Experimental distributions of Peak 1 for three patterns per cell. Conventions as in (A). E. Experimental heatmap for Peak 1 differences during excitation. Conventions as in (B). F. Simulation heatmap, analogous to (E), for Peak 1 during excitation. (G-I) STPR during Inhibition. G. Experimental distributions of STPR for three patterns per cell. Conventions as in (A). H. Experimental heatmap for STPR differences during inhibition. Conventions as in (B). I. Simulation heatmap, analogous to (H), for STPR during inhibition. (J-L) STPR during Excitation. J. Experimental distributions of STPR for three patterns per cell. Note that patterns with heterogeneous initial synaptic responses (Peak 1) do not always show heterogeneity in STPR. Conventions as in (A). K. Experimental heatmap for STPR differences during excitation. Conventions as in (B). L. Simulation heatmap, analogous to (K), for STPR during excitation. (M-N) Summary of Pattern Heterogeneity. M. Experimental data: Percentage of cells showing any significant (p < 0.05) difference between pattern responses for Peak 1 and STPR (Inhibitory & Excitatory conditions). N. Simulation data: Analogous summary to (M).

The simulations provide an insight into the source of this heterogeneity, since they allow us to separate the contributions of synaptic weight and connectivity distributions. GluR excitatory conductances were 0.807±0.223 nS (mean±standard deviation) whereas there was a uniform GABAR conductance of 10.053 nS. However, there was a single stage of connection from CA3 to CA1 (probability of connection p=0.02) but two stages between CA3 to Interneurons (p=0.01) and interneurons to CA1 (p=0.01). Thus we infer that the major source of heterogeneity in E(I)PSCs is because sparse connectivity leads to different numbers of active synapses for different patterns.

### Synaptic stochasticity accounts for most trial to trial variability

We next zoomed in to the single-synapse level, to examine how stochasticity in synaptic release might account for the trial-to-trial variability of synaptic responses. To do this, we used the 12 trial repeats recorded for each pattern. While there is no absolute mapping between number of stimulated squares and number of activated synapses due to expression differences, we expect that 1 square patterns should have more release failures and 5 and 15 square patterns should exhibit fewer response failures. Our observations were consistent with this both in experimental and simulated data (Figure 5A-D).

**Figure 5:**
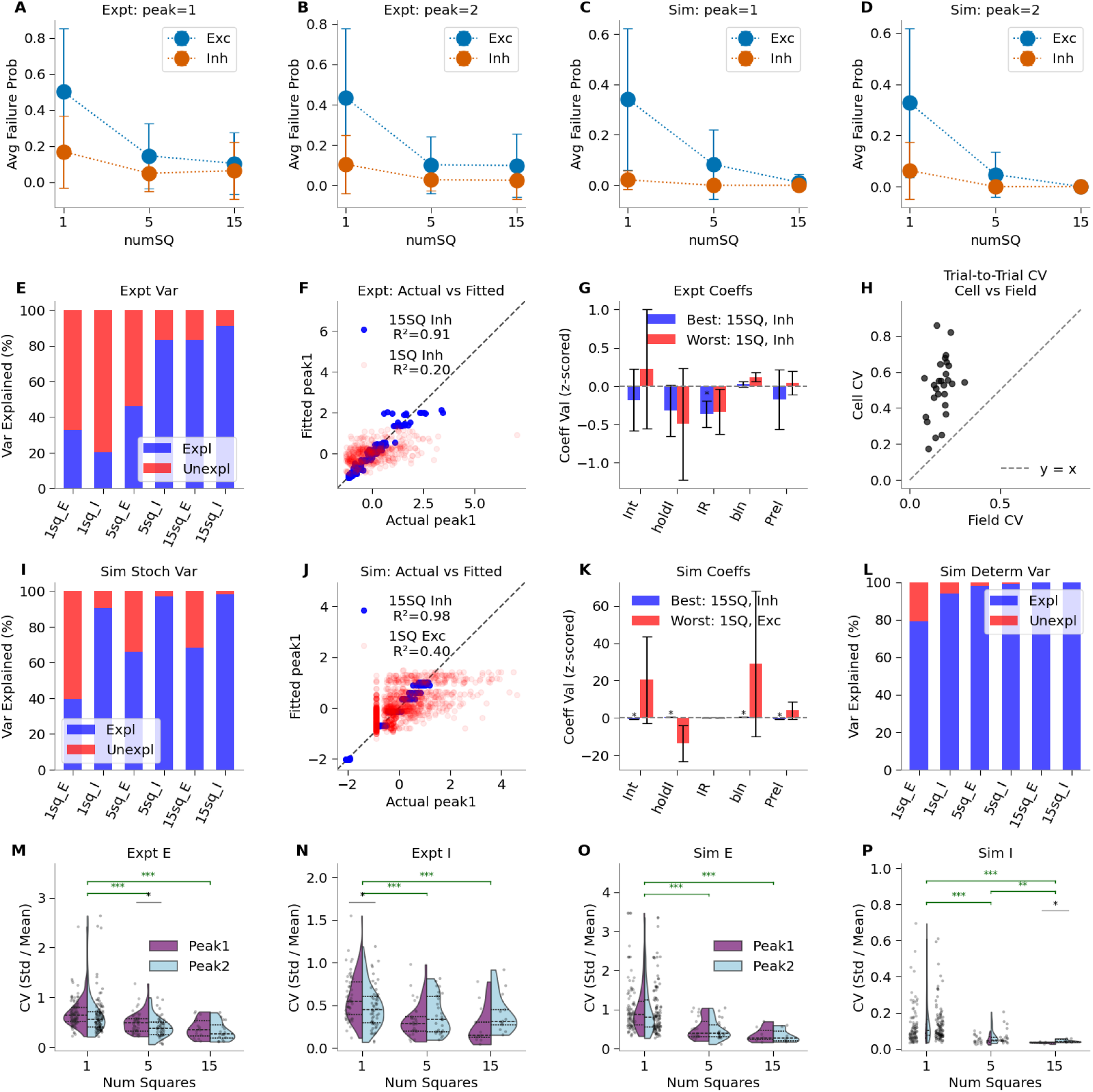
Synaptic stochasticity accounts for most trial-to-trial variability. (A-D) Average failure probability (± SEM across cells) for Peak 1 and Peak 2 responses as a function of the number of stimulated squares (1, 5, 15). Data are shown for simulated (Sim) and experimental (Expt), separated by excitatory (Exc, blue) and inhibitory (Inh, orange) conditions. A. Experimental data: Failure probability for P1 responses. B. Experimental data: Failure probability for P2 responses. C. Simulation data: Failure probability for P1 responses. D. Simulation data: Failure probability for P2 responses. (E-L) Generalised Linear Model (GLM) Analysis of Peak 1 Variance. GLMs were used to determine how much of the trial-to-trial variance in normalized Peak 1 amplitude could be explained by categorical variables (cell ID, pattern ID) and continuous predictors (holding current, input resistance, baseline noise (std. dev.), and a model of probability of release based on number of squares). E. Experimental Data: Variance Explained. Stacked bar chart showing the percentage of Peak 1 variance explained by the GLM (blue) and unexplained (residual, red) across different pattern sizes (1, 5, 15) and clamp potentials (E: Excitatory, I: Inhibitory). F. Experimental Data: Actual vs. Fitted Peak 1. Scatter plot of actual normalized Peak 1 amplitudes against GLM-fitted values for the experimental conditions showing the best and worst goodness-of-fit. The identity line (y=x) is shown in black. G. Experimental Data: GLM Coefficients. Bar chart comparing the normalized GLM coefficients for physiological predictors (Int: Intercept, holdI: Holding Current, IR: Input Resistance, bln: Baseline noise (std. dev.), Prel: model) for the best-fit and worst-fit parameters identified in (F). H. Experimental Data: Trial-to-Trial CV Comparison (Cell vs. Field). Scatter plot comparing the coefficient of variation (CV) of cellular Peak 1 responses to the CV of simultaneously recorded field potentials for excitatory cells (n=3), illustrating correlated trial-to-trial variability. The identity line (y=x) is shown. I. Stochastic Simulation Data: Variance Explained. Similar to (E), but for data from stochastic simulations. J. Stochastic Simulation Data: Actual vs. Fitted Peak 1. Similar to (F), but for stochastic simulation data. K. Stochastic Simulation Data: GLM Coefficients. Similar to (G), but for stochastic simulation data. L. Deterministic Simulation Data: Variance Explained. Similar to (E) and (I), but for data from deterministic simulations (single trial per pattern) to highlight variance explained by intrinsic differences between cells and patterns. (M-P) Coefficient of Variation (CV) of Peak Amplitudes. Split violin plots showing the distribution of CV (Std/Mean) for Peak 1 (P1, purple) and Peak 2 (P2, sky blue) responses across different numbers of stimulated squares (1, 5, 15). Individual data points (CV per pattern per cell) are overlaid. Horizontal lines with asterisks indicate significant differences between P1 and P2 CV distributions within a 1, 5, 15-square patterns (t-test: * <0.05, ** <0.01, *** < 0.001). Lines connecting numSQ groups with asterisks indicate significant differences in CV(P1) between 1, 5, 15-square patterns. M. Experimental data: CV comparisons for excitation. N. Experimental data: CV comparisons for inhibition. O. Simulation data: CV comparisons for excitation. P. Simulation data: CV comparisons for inhibition.

We then investigated the sources of trial-to-trial variability using a generalized linear model (GLM) incorporating cell physiological parameters, instrument noise, and synaptic release probability. For the experimental dataset, the GLM’s ability to account for response variance increased with stimulus size: from 1-square, to 5-square, to 15-square patterns, where nearly all variability was explained (Figure 5E).

The simulations afforded us the ability to separate out two contributions to inter-trial variability: experimental biological and equipment noise, and the contribution of synaptic release stochasticity. In the simulated dataset, for distinct cells we used different pattern density terms as in Figure 2, corresponding to higher ChR2 expression. We also used distinct random number seeds to set up the connectivity for each cell. Together these formed the basis for categorical variation between simulated cells. We first ran the simulations with stochastic calculations for the synapses, and found that the explained variance for EPSCs was comparable with experiment for each of the 1, 5 and 15 square patterns (Figure 5I-K). We infer that stochasticity accounts for most of the unexplained experimental variance in EPSCs. Surprisingly, for the IPSCs, there is very little unexplained variability in the simulations. We speculate that cellular noise may account for more of the experimental variance for IPSCs since the cell is held at 0 mV and is consequently stressed. As a final test, we reran the simulations with the stochastic calculations replaced by deterministic ones (Figure 5L). Here there was very little unexplained variability in all cases, further reinforcing the primary role of stochasticity in the variance.

We then examined the coefficient of variation (CV) between trials as another readout of variability. As expected, larger stimuli show reduced coefficient of variation (CV). We do not find significant differences in CV from peak 1 and peak 2 responses, suggesting that the relative reliability of synaptic release did not change during STP under our experimental conditions (Figure 5M-P).

Thus we were able to show that probabilistic (stochastic) synaptic release accounted for most of the features of inter-trial variability, particularly for excitatory synapses.

### Cell physiology and stimulus variability do not explain trial to trial STP variability

Having characterized E(I)PSC variability, we next looked at trial to trial variability in STP for repeats of each stimulus pattern. As before, the STP ratio (STPR) was calculated as the difference in peak amplitudes normalised to the sum of peaks. We found that patterned stimulation results in a wide range of STP profiles in CA1 cells both in excitation and inhibition on a trial to trial basis that was not explained by cell physiology and stimulus variability (Figure 6A-C shows a representative cell). The stochastic simulations qualitatively matched these distributions (Figure 6D-F shows a representative simulated cell). Field response variability was not correlated with cell response variability in either excitation (Figure 6G) or inhibition (Figure 6H). We then examined the relationship between cell physiological properties and STPR using a generalized linear model (GLM) analysis. Experimental cell physiological parameters, baseline noise and ChR2 desensitization revealed poor predictive power for these intrinsic and experimental sources of variability (Figure 6I). In the case of simulations, as explained above, the cell differences were due to input strength and connectivity properties. Again, these did not explain most of the variability in STPR.

**Figure 6:**
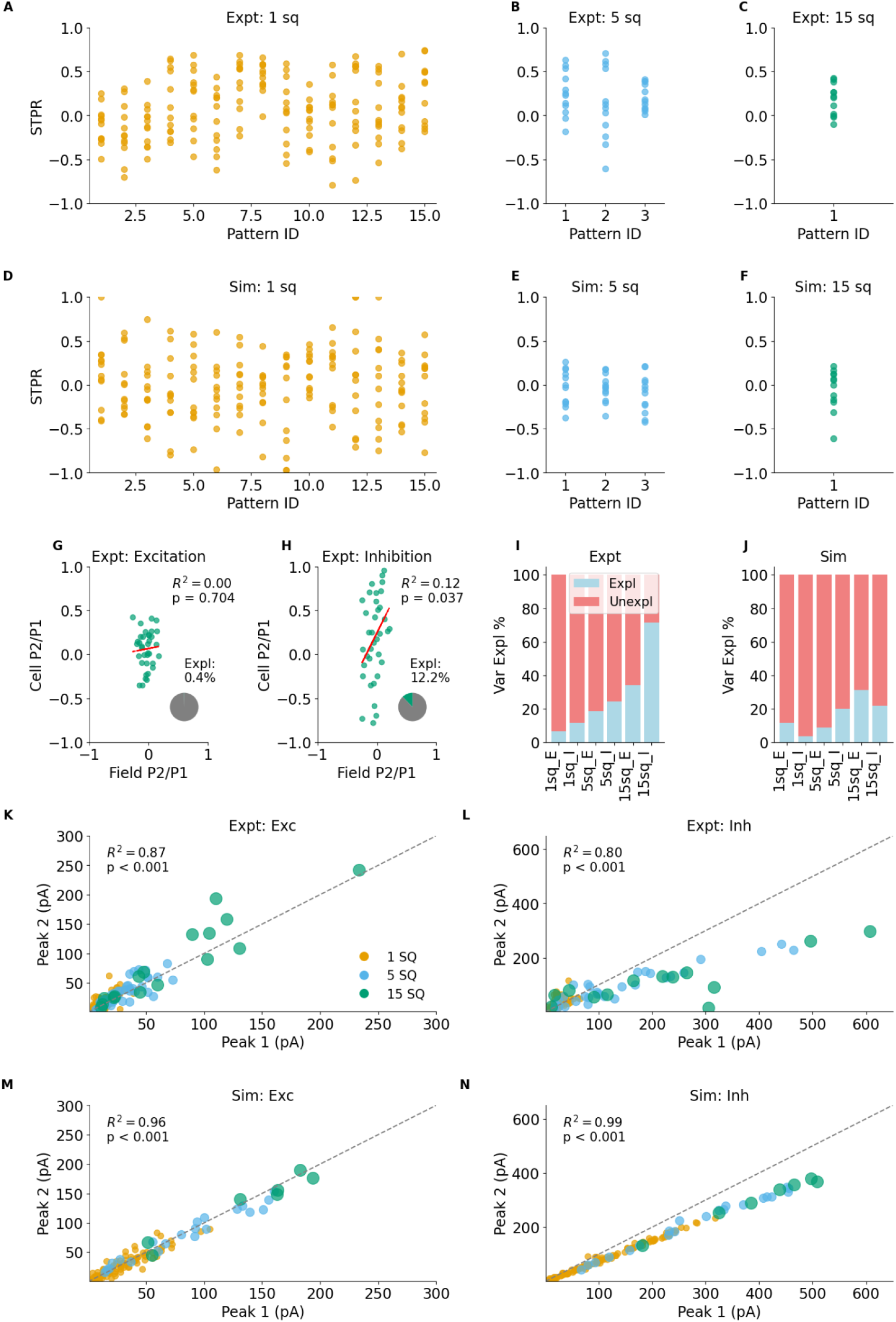
Model captures experimental STP dynamics. (A-C) The STPR is shown for a single representative experimental neuron for excitation. Each panel corresponds to a different stimulus strength: 1 sq (A), 5 sq (B), and 15 sq (C). Points represent the STPR for individual patterns. (D-F) Corresponding STPR plots for a representative simulated neuron. (G-H) Correlation between cellular and field potential STPR for the 15-sq stimulus in experimental data, for excitation (G) and inhibition (H). The red dashed line is the linear fit; R^2^ and p-value are shown. A simple linear regression (Ordinary Least Squares) was performed to model the cellular STPR as a function of the field STPR. The R-squared (R2) value is shown to quantify the proportion of variance in the cellular STPR that is predictable from the field STPR. The inset pie chart directly visualizes this R2 value: the green slice represents the percentage of variance explained by the model (R^2^*100), while the larger gray slice represents the unexplained variance. The small size of the green slice demonstrates that the field potential is a poor predictor of the STPR of an individual neuron. (I-J) Percentage of STPR variance explained by a Generalized Linear Model (GLM) for experimental (I) and simulation (J) data. The GLM includes cell physiological parameters (holdI, IR), baseline noise (bln_std_cell), and stimulus strength (P_release) as predictors, while also factoring in cellID and patternID as categorical variables. Bars show explained (light blue) vs. unexplained (light coral) variance for each stimulus strength and clamp condition (E/I). (K-N) Peak amplitude of the second response (P2) vs. the first (P1). The grey dashed line is the unity line. Points are colored by stimulus strength. (K-L) Experimental data for excitation (K) and inhibition (L). (M-N) Corresponding simulation data showing similar trends.

We finally performed direct trail-wise comparisons between peak 2 and peak 1 as another readout of STP. Excitatory synapses showed little or no STP (Figure 6K) whereas inhibitory synapses show strong short term depression (Figure 6L). By design, the simulations matched these closely (Figure 6M, N).

### E and I contributions were validated under physiological conditions

To validate the contributions of E and I inputs and the resultant STP profiles under physiological conditions, we performed current clamp experiments using a single 15-square pattern stimulation. Each trial consisted of a probe pulse followed by an 8-pulse train delivered at 20 Hz, with the probe preceding the train by 1 second. We recorded responses in the presence (Figure 7A) and absence of GABA_A receptor-mediated inhibition (Figure 7B). Upon bath application of gabazine (20 µM), we observed a robust increase in PSP amplitude (Figure 7B, D), consistent with disinhibition of CA1. We recapitulated key features of CA1 STP including response waveform and STP using simulation (Figure 7B, D). We were able to replicate the classic STP train of initial facilitation followed by depression in experimental (Figure 7F) data. Here again we varied the pattern density parameter in our current clamp model to account for the ChR2 expression level differences between recorded cells. Using this approach, we were able to match the distributions of peak 1 amplitudes (Figure 7I, K) as well as STP ratios (Figure 7J, L) with experiments. To assess cell to cell variability in STPR, we performed bootstrap resampling of STP ratios for the initial paired pulse in the train across all possible cell combinations and obtained p-values for cell similarity. We then performed hierarchical clustering of cells from the resulting p-value matrix and found some degree of cell clustering based on their STP profiles. STP does not have a significant dependence on initial release, and there is considerable scatter, especially at low release values (R2=0.09, p=0.46) (Figure 7N). This is similar to our observations of nearly flat STPR with Peak 1 from voltage clamp (Figure 2M). This trend is weakly negative in simulated data (R2=0.32, p=0.002) (Figure 7P). We also assessed synaptic summation by dividing the 15-square stimulus into three random 5-square patterns, presented separately to measure individual responses. The observed response to the full 15-square pattern was then compared to the linear sum of the 5-square responses. We found predominantly sublinear summation, with maximal summation at the second peak of the train (Supplementary Figure 4), consistent with (Bhatia et al., 2019) and (Asopa & Bhalla, 2025). Thus our recordings and simulations in more physiological current clamp conditions were substantially consistent with the earlier voltage clamp readings, and with previous work.

**Figure 7:**
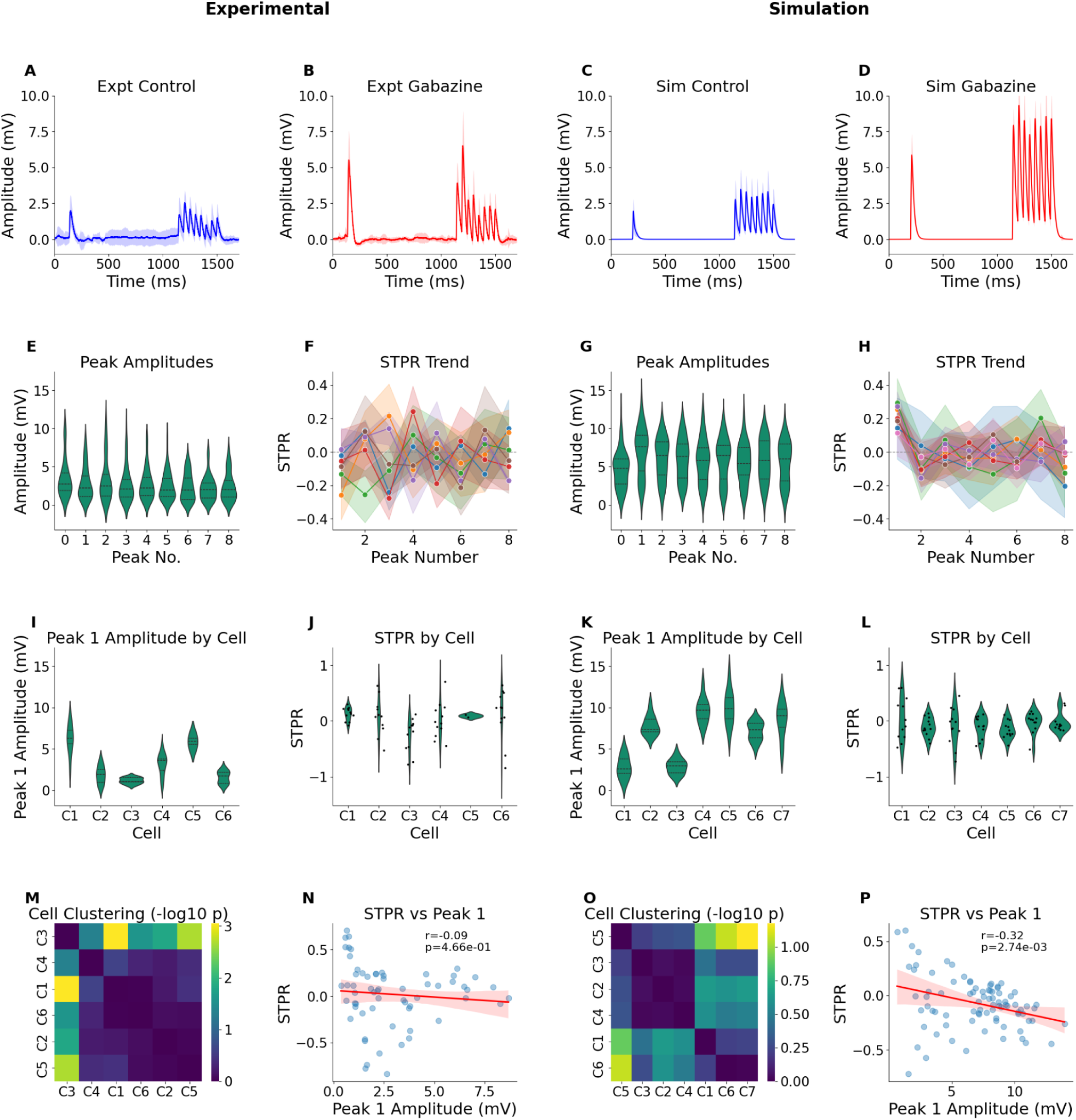
Model captures experimental STP dynamics under physiological conditions. (A-D) Raw Traces in response to 1 test pulse followed by a burst of 8 pulses, from representative neurons. Individual trials are shown as faint black lines, with the mean trace overlaid in blue (control) or red (gabazine). Experiment, control (A); Experiment, Gabazine (B); Simulation, control (C); Simulation, GABA weights set to zero to simulate Gabazine (D). The model cell membrane resistivity RM was reduced from 1.0 to 0.3 S/m^2^ to account for the much smaller dendritic extent and hence membrane area of the simulated neuron. (E, G) Peak Amplitude Distribution for experiment and simulation respectively for probe and eight pulse train across all cells and trials. (F, H) STPR by Cell for experiment and simulation. Violin plots showing the distribution of the STP ratio for the first two peaks of each individual cell. Black dots represent individual trials. (I, K) Peak 1 Amplitude by Cell for experiment and simulation. Violin plots showing the distribution of the first peak’s amplitude (peak 1) for each individual cell. (J, L) STPR Dynamics for experiment and simulation. Line plots tracking the STPR value across subsequent peaks (peak0 is the probe peak). Each colored line represents a different cell. (M, O) Cell Clustering for experiment and simulation based on their STPR profiles. Heatmaps show the statistical similarity between cells . Color intensity represents the -log10(p-value), with cells ordered by hierarchical clustering. (N, P) Correlation between the first peak’s amplitude (peak1) and the STP ratio of the first two peaks of the 8-pulse train, with a linear regression line. Both experiment and simulation are quite scattered and nearly flat, but simulation has a small negative slope.

### Synaptic heterogeneity and stochasticity have diverse effects on signal transformations

Having tuned our model to experimental readouts of heterogeneity and other forms of variability, we finally deployed the model to examine the contributions of various mechanisms to the transformation of an autocorrelated input signal. Biologically detailed models are uniquely well-suited to this analysis, because one can isolate terms such as stochasticity, heterogeneity in connections, and STP relationships in a manner which is not accessible experimentally. We generated autocorrelated input patterns with mean frequencies of 25 to 250 Hz (methods, Figure 8A). We delivered these patterns to our simulated optical input grid for 1, 5 and 15 square patterns, each with a pattern density of four (methods). The reference model with stochasticity and synaptic heterogeneity completely decorrelated the weaker 1-square input signal (Figure 8Bi, Bii, Bv) but at 15 squares the stimuli were strong enough to cause the postsynaptic cell to follow low-frequency input, hence retaining some autocorrelation (Figure 8Biii, Biv, Bv). Thus the reference model mostly leads to decorrelation, but strong low-frequency input retains correlation.

**Figure 8:**
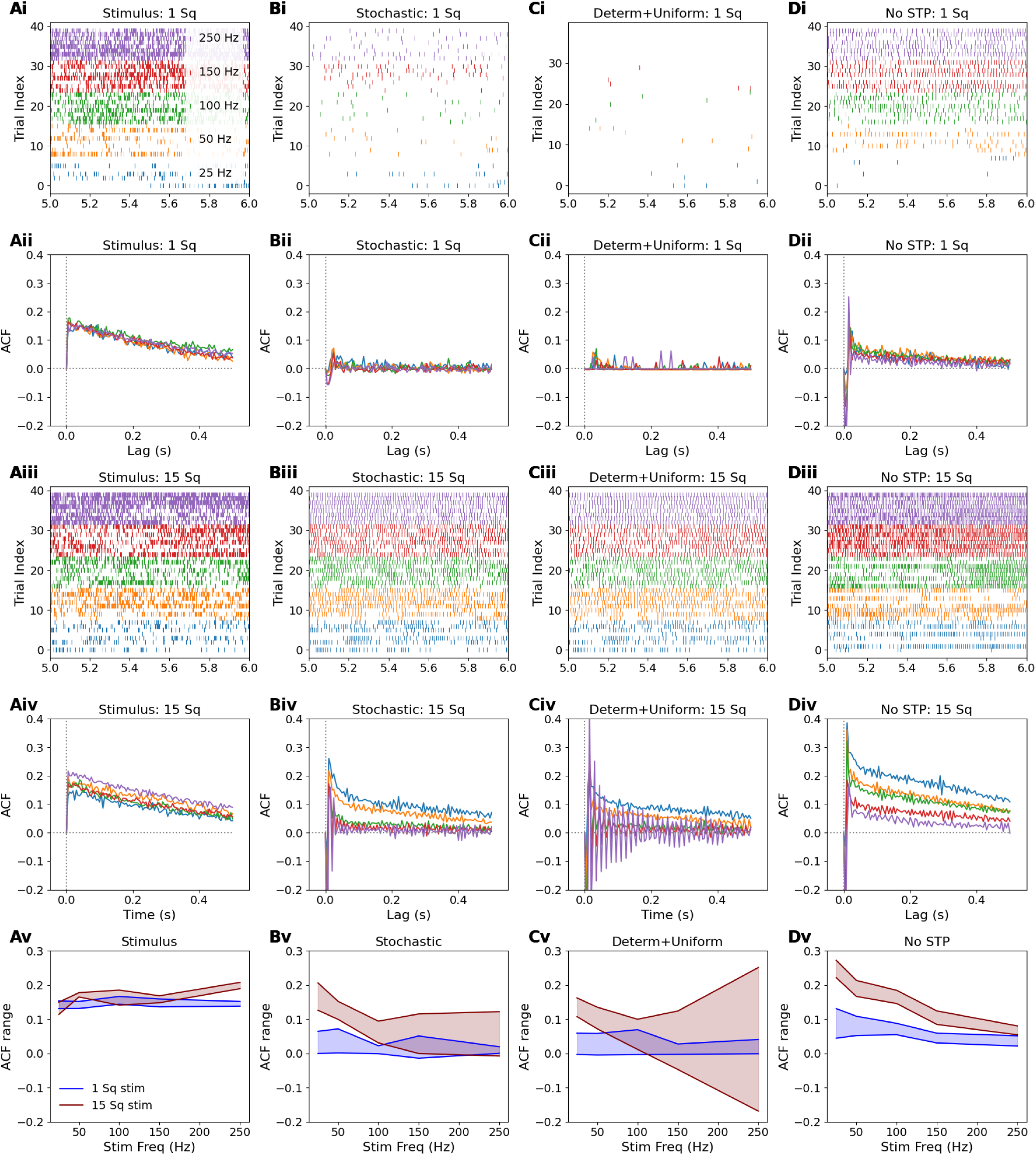
Effect of stochasticity and heterogeneity on output decorrelation. A. Stimulus. Ai: 1-second sample raster of pseudo-random autocorrelated input spike pattern. Different colors indicate different mean frequencies; same colors apply for all plots. Aii: Autocorrelation function (ACF) of the input pattern. Aiii and Aiv: equivalent to Ai and Aii respectively, different seed for 15-square runs. Av: stimulus autocorrelation is nearly independent of frequency. Subsequent runs in B, C and D used the same random number seeds and hence the exact same pseudo-random stimulus spike trains as reported here for A. B. Responses for reference stochastic + heterogeneous model. Bi: Output raster, 1 square stimulus. Bii: Complete decorrelation occurs for 1-square inputs. Biii: Output raster, 15 squares. Biv: Selective decorrelation for inputs above 50 Hz, for 15-square inputs. Bv: There is strong decorrelation except low-frequency 15-square. C. Output spiking patterns and ACFs for deterministic + uniform model. Ci: spiking is low for 1 square. Cii: ACF is flat. Ciii: Spike raster 15 square. Civ: The ACF reports strong periodicity for 150 and 250 Hz inputs which is regular out to almost 200 ms. Cv: Summary plot shows decorrelation for 1 square, but increasing oscillations (positive and negative bounds) at 150 and 250 Hz. D. Responses for model with non-STP synapses, but with stochasticity and heterogeneity retained. Di, Dii: spiking and ACF show decorrelation for 1-square input. Diii, Div: frequency-dependent decorrelation, where the cell amplifies the correlation of low-frequency inputs, but reduces it for high-frequencies. Dv: Decorrelation increases with frequency for 15-square inputs.

We next eliminated all forms of variability. We programmatically changed all the stochastic synapses to deterministic, and made all the connection weights uniform. This led to very weak firing for the 1-square input (Figure 8Ci, Cii). In striking contrast to the reference model, the 15-square inputs led to strongly periodic activity (oscillatory autocorrelation function) at higher frequencies of input (Figure 8Ciii, Civ, Cv). We speculate that under these conditions all the synapses contribute to a slow averaging of inputs, leading to nearly uniform firing. Weaker periodicity also emerged if just stochasticity was eliminated (Supplementary Figure 5Div). Thus the presence of stochasticity and heterogeneity rescues the neuron from periodic activity.

Third, we asked what happens if STP itself is eliminated both from E and I synapses, while retaining the same stochasticity and heterogeneity of the reference model. As before, low-strength inputs lead to decorrelation (Figure 8Di, Dii). High-strength (15-square) inputs lead to a clear segregation of output autocorrelation levels by frequency (Figure 8Diii, Div). Remarkably, at low frequencies the output autocorrelation was even higher than that of the input (Figure 8 Div, Dv). Contrasting this with the reference model, we find that STP increases decorrelation, and without STP the neuron has increasing decorrelation for higher-frequency inputs (Figure 8Bv vs Dv).

We also examined what happens if EI balance is perturbed, by programmatically setting all GABA weights to zero and lowering Glu weights to compensate. The outputs did not differ much from the reference model (Supplementary material). Thus our model predicts that EI balance has little effect on input-output decorrelation.

Overall, we find that reference levels of heterogeneity, stochasticity and STP yield decorrelation, that uniform and deterministic synapses may give periodic firing, and that STP flattens out the frequency-dependence of decorrelation.

## Discussion

We used optical spatiotemporal patterned inputs to the CA3-CA1 circuit to characterize different contributions to heterogeneity and stochasticity in input-output transforms and in short-term plasticity. Heterogeneity between neurons and between patterns composed of different groupings of light spots were well explained by differences in ChR2 expression and connectivity. Variability between repeated trials was substantially explained by stochastic synaptic release. The distribution of summation responses was also consistent with these major contributors to heterogeneity and variability. At each stage we demonstrated that a multiscale model of stochastic chemistry at synapses, neuronal physiology, and network connectivity could capture most of the observations. Finally, we used the model to examine how heterogeneity and variability contribute to decorrelation of spiking output, as opposed to periodic or frequency-dependent autocorrelated output spike trains.

### Heterogeneity across scales

Individual synapses: Our findings are in line with a substantial body of literature on the diversity of synaptic properties, which play a crucial role in enhancing the richness and flexibility of neuronal and network computations. At the level of single synapses, Nusser et al showed that the AMPA receptor content on CA3->CA1 synapses ranged from <3 to 90 (Nusser et al., 1998). Hippocampal GABAergic synapses are also highly heterogeneous (Aradi et al., 2002; Biró et al., 2006). Sasaki et al. used paired CA3-CA1 recordings to show that action potential transmission in organotypic hippocampal slices was reliable but that each synapse behaves independently, stochastically, and with highly heterogeneous postsynaptic effects (Sasaki et al., 2012). Each of these features is present in our data and model.

Single synaptic STP properties are also diverse in thalamocortical (Díaz-Quesada et al., 2014b) and barrel cortex synapses (Buchholz et al., 2023b). Debanne et al. (Debanne et al., 1999) measured LTP as well as STP in rat hippocampal slice culture and compared multi-unitary and unitary EPSPs to show heterogeneity of STP as well as LTP. Our study also finds heterogeneous STP, but enables a more circuit-level analysis as we simultaneously obtained E and I synaptic weights and STP properties on distinct projections converging onto CA1 neurons in the hippocampus. One caveat is that our inhibitory input readouts are a composite of synaptic transfer functions from CA3 to interneurons and then from interneurons to the CA1 cell, as stressed in figures 3 and 4.

A distinguishing feature of our recordings is that we measure functional convergence of well-defined, but sparse input patterns. Many studies employ field electrode stimulation to measure synaptic responses (Sun et al., 2018b), but this leads to massive activation of the Schaffer collaterals and possible saturation of the E or I pathways (Supplementary Figure 1). Finer granularity can be obtained by using arrays of stimulus electrodes (Parameshwaran & Bhalla, 2012) and by reducing stimulus strength towards minimal stimuli (Dobrunz & Stevens, 1997b), but the triggering of identified axons by such methods is uncertain and to our knowledge arrays of such stimuli have not been utilized. At the most specific level, a few studies have performed dual-patch recordings from CA3 and CA1 pyramidal neurons (e.g., Debanne et al., 1999; Sasaki et al., 2012) but these also have not looked at convergence of multiple inputs. The Blue Brain Project has performed simultaneous multi-patch recordings including STP analysis in the somatosensory cortex (Markram et al., 2015) but to our knowledge this has not been done in the hippocampus. Thus our analysis of heterogeneity in E(I)PSCs and STP responses to multi-square pattern responses (Figures 3, 4 and 5) uncovers further details about the distribution of responses to be expected *in vivo*.

There are many modelling studies of the hippocampus at various levels of detail. Traub and Jeffries implemented one of the first biophysically detailed network-level models of epileptiform activity in CA3 (Traub & Jefferys, 1994). Hippocampal memory models have also been developed (Cutsuridis et al., 2010). There are recent resources providing connectivity and biophysics for large-scale hippocampal models (Ecker et al., 2020; Gandolfi et al., 2023). Such models function at whole-network scales. At a cellular level of detail, there have been several data-driven models of CA1 pyramidal neurons (Gasparini et al., 2004). At a still finer level, the classic studies of presynaptic STP (Tsodyks & Markram, 1997) have led to a family of abstract STP models (Hennig, 2013), as well as more biologically motivated models for heterogeneity in release: (Hanse & Gustafsson, 2001; Neher, 2015). At the other extreme, STP has been modeled at single-molecule reaction-diffusion levels of detail (Nadkarni et al., 2010, 2012). We have recently implemented a model of the hippocampal circuit that spans all these levels (Asopa & Bhalla, 2025) and the current study fine-tunes this model for the study of heterogeneity in the circuit. To our knowledge there are no other studies that integrate all these scales. A primary consideration in our model development was the ability to simulate many trials and stimulus combinations, amounting to several hours of electrophysiological recording time, yet to do so at a level of detail that matched all aspects of our experiments. Thus we have chosen abstract neuron models for CA3 and interneuron arrays, a reduced multicompartment conductance based electrophysiological model for CA1, and Gillespie-algorithm (Gillespie, 1977a) based stochastic chemical reaction kinetics for the STP. Our model is open source and lightweight, and runs about 20x slower than real-time on a laptop, thus providing a readily reusable resource for further exploration of this system.

Heterogeneity in STP is not merely noise but an important functional feature. The variability in STP profiles across different CA1 synapses allows for a more nuanced and adaptive response to incoming stimuli shaping the learning capacity of neural networks (Tsodyks & Markram, 1997). Several of the features we examine have been considered using a more abstract mathematical analysis based on Tsodyks STP formalism (Rosenbaum et al., 2013). Our findings are in partial accord with their predictions, as follows: We have both STP (Glu synapses) and STD (GABA synapses) in our model, and we find that together these forms of plasticity give rise to the decorrelation of input trains. This matches somewhat with their findings of STD-dependent decorrelation. However, we find that going from 1 to 15 square inputs (more synapses) reduces correlation, unlike their prediction of more correlation. Our study agrees that faster input firing gives more decorrelation, but quite notably we find that deterministic inputs give rise to periodic firing especially at high-frequency inputs, in contrast to their result of decorrelated inputs. We suggest that this means that the heterogenous, stochastic system is less likely to fall into degenerate simple periodic activity patterns. We suggest that our more biologically detailed model has significantly richer dynamics, leading to a wider range of emergent properties. At a broader level our findings agree with and substantially extend previous work which supports the importance of cellular and circuit heterogeneity, plasticity, and stochasticity in generating interesting transformations of input signals.

## Methods

### Animals

Transgenic mice C57BL/6-Tg(Grik4-cre)G32-4Stl/JNcbs with tissue-specific expression of Cre recombinase in the CA3 region of the hippocampus were used. Cre recombinase expression is driven by the endogenous promoter/enhancer elements of the glutamate receptor, ionotropic, kainate 4 (Grik4) gene (source: Jackson Laboratory, Strain ID: 006474). All animals were obtained from Jax and bred in NCBS Animal Care and Resource Center (ACRC) in Specific Pathogen Free (SPF2) conditions. The breeding scheme involved a male hemizygote (m) crossed with a female non carrier (f). Animals were maintained with a light and dark cycle of 14:10 (14 hour light: 10 hour dark) and *ad libitum* access to food and water. All animal procedures were operated and approved according to the National Centre for Biological Sciences Institutional Animal Ethics Committee (NCBS IAEC) guidelines under IAEC Project# NCBS-IAE-2022/8 (R2ME).

### Viral transfection of Channel Rhodopsin protein using Cre-Lox

Cre-positive animals were identified by genotyping and used for optogenetic experiments. 25–35 days old transgenic mice were injected with AAV-FLEX-rev-ChR2-tdtomato (Plasmid# 18917) virus to achieve Cre-dependent viral expression of ChR2-tdTomato in CA3 pyramidal cells. The virus was diluted 1-5x in sterile PBS to a titer of 1-5 x 10^12 GC/ml and stored at -80°C. Stereotaxic injections were made into the hippocampal CA3 region using the following coordinates: -2.0 mm RC, ±1.9 mm ML, -1.5 mm DV. ∼700 nl solution was injected into the CA3 region of the left or right hemisphere with brief pressure pulses of 5-15 psi, 10-20 ms pulses using Picospritzer-III (Parker-Hannifin, Cleveland, OH). Animals were allowed to recover for at least four weeks following surgery. AAV-FLEX-rev-ChR2-tdtomato was a gift from Scott Sternson (Addgene plasmid# 18917; RRID:Addgene_18917). Injected animals were transferred to a separate holding facility until experiments.

### Optogenetic stimulation

Optogenetic stimulation was delivered using a Polygon400 Digital Mirror Device (Polygon 400G from Mightex Pvt. Ltd, Ontario). The Polygon DMD device was mounted into the infinity-path of an Olympus BX61WI microscope and features a micro-mirror array of 684X608. During optical stimulation the water immersion 40X objective is centered on the CA3 cell body layer. The light source had a total estimated maximum intensity of 6.7 mW and each spot of the 24X24 optical grid had an absolute power of approximately 14.47 µW and a power density of 100 mW/mm2 (Asopa & Bhalla, 2025). Optical grid images (in png or bmp) were generated using custom python code and uploaded to Polygon using Polyscan2 (Mightex) and protocols were written using Clampex (Molecular Devices). During multi-square stimulation, adjacent squares were avoided to minimize overlap. The light intensity was kept at 100% with a pulse width to 5 ms for all experiments. The intersweep interval was set to 1-square stimulation and 10 seconds for 5- and 15-square stimulation to prevent ChR2 desensitization and long term plasticity effects.

### Slice preparation

8–12 weeks (4–8 weeks post virus injection) old mice were anesthetized with isoflurane and decapitated following cervical dislocation. The brain was immediately transferred to ice-cold cutting solution (87mM NaCl, 2.5mM KCl, 7mM MgCl2.6H2O, 0.5mM CaCl2.2H2O, 25mM NaHCO3, 1.25mM NaH2PO4.H2O, 75mM Sucrose). Hippocampus from the hemisphere where virus injection was done was dissected out and 400 um transverse slices were prepared using a vibratome (Leica VT1200). Slices are cut in ice-cold ACSF containing (87mM NaCl, 2.5mM KCl, 7mM MgCl2.6H2O, 0.5mM CaCl2.2H2O, 25mM NaHCO3, 1.25mM NaH2PO4.H2O, 75mM Sucrose) while carbogen was continuously perfused through the buffer tray of the vibratome. Slices were transferred using a pasteur pipette into recording aCSF (124mM NaCl, 2.7mM KCl, 1.3mM MgCl2.6H2O, 2mM CaCl2.2H2O, 26mM NaHCO3, 1.25mM NaH2PO4, 10mM D-(+)-glucose, pH 7.3-7.4 and osmolarity of 305-315 mOsm) at room temperature onto a nylon mesh tray kept inside a beaker that is continuously perfused using carbogen (95% O2, 5% CO2). Slices were incubated in the dark for one hour before being transferred to a slice chamber on an inverted Olympus microscope stage, where they were held in place with a slice holder (Warner Instruments WI 64-0246). Recording aCSF that was continuously perfused using carbogen was heated to physiological temperatures (32-34°C) using an inline heater (Warner Instruments TC-324B) and ran through the slice chamber at 1 ml/sec.

### Electrophysiology

Whole cell recording pipettes of 4-6 MO were pulled from thick-walled borosilicate glass capillaries from Sutter Instruments (1.50mm O.D., 0.86mm I.D., 1Ocm long) on a P-97 Flaming/Brown micropipette puller (Sutter Instrument, Novato, CA). For current clamp experiments, pipettes are filled with internal solution containing (in mM): 130 K-gluconate, 5 NaCl, 10 HEPES, 1 EGTA, 2 MgCl2.6H2O, 2 Mg-ATP, 0.5 Na-GTP and 10 Phosphocreatine, pH 7.3 - 7.4, osmolarity 290 - 295 mOsm. For voltage clamp experiments, instead of K-gluconate, 130 mM Cs-gluconate was used (from Gluconic Acid + CsOH). Cells were visualized using infrared microscopy and differential interference contrast (DIC) optics on an upright Olympus BX61WI microscope (Olympus, Japan) fitted with a 40X (Olympus LUMPLFLN, 40XW), 0.8NA water immersion objective. Recordings were acquired using Axon amplifier 700B and Digitizer 1400. Electrical recordings were acquired at 20 kHz and filtered at 10 kHz using Axon Amplifier 700B and Digitizer 1400 (Molecular Devices, San Jose, CA). Clampex and pCLAMP softwares (Molecular Devices) were used for protocol writing and data acquisition. For field recordings, a recording electrode was kept next to the CA3 cell body layer to measure the net optogenetic activation of CA3.

For the voltage clamp experiments, cells were held at either -70 or 0 mV. Cells were included in the analysis if their input resistance varied by less than 25% over the course of the recording. For these cells, the mean ± standard deviation of input resistance fluctuation was 9.82 ± 5.70%. The baseline holding current showed a mean percent fluctuation of 18.78 ± 12.05% and an absolute fluctuation of 15.78 ± 9.13 pA. For the current clamp experiments, cells were held at their resting membrane potential (RMP: Mean=-66.81 mV, Std Dev=6.17 mV; Input Res: Mean=71.44 MOhm, Std Dev=64.15 MOhm). Cells were eliminated if holding current went above 100 pA or fluctuated more than 20 pA during the course of the experiment. Cells were eliminated from analysis if resting membrane potential fluctuated more than 5 mV during the course of the experiment (Std Dev=3.84 ± 3.01 mV; CV=0.0582±0.0450). To isolate the effects of inhibition, a selective and competitive GABA_A_ receptor antagonist Gabazine (HelloBio HB0901 - SR 95531) was used at 20 uM concentration. Recordings with Gabazine were made after complete perfusion was ensured, using holding current stability as a proxy.

### Model

Our multiscale model was almost identical to (Asopa & Bhalla, 2025) with the exception of the choice of parameters for the chemical signaling model, which were tuned to the current dataset. The main outcome of this tuning was to have steeper STD on the GABA synapses, as per Figure 2K. The reaction scheme for both Glu and GABA release was identical (Figure 2G). The neurotransmitter release step was coupled to an alpha-function conductance-based AMPA receptor model on the spine (Glu release), and an alpha-function model for GABA on the dendrite. There were 100 spines, each with a presynaptic bouton incorporating a distinct instance of the presynaptic reactions, and 200 GABA boutons, again each modeled independently. The CA1 cell model was a ball-and-stick model with a 10-micron soma and a 2 micron diameter x 200 micron length dendrite, on which the respective synapses were evenly distributed (Figure 2H). Na and K_DR channels were present at the soma, at subthreshold levels for all simulations except the spiking models in figure 8 (Supplementary Data 2, Table 5).

The CA3 was modeled as a 16x16 array of single compartments (Figure 2I) whose membrane potential was set above threshold if the simulated optical pattern at that location was ‘on’. Active cells triggered presynaptic signaling on the Glu boutons.

The interneurons were also modeled as a 16x16 array of thresholded single compartments, whose potential was assigned above threshold if any of the CA3 inputs was ‘on’. Interneuron output was sent to the GABA boutons on the CA1 neuron model.

Connections were implemented as a 256x100 matrix from CA3 to CA1 Glu synapses (connection probability p = 0.02), 256x256 matrix from CA3 to interneurons (p = 0.01), and 256x200 matrix from interneurons to CA1 GABA synapses (p = 0.01).

Stimuli were delivered as 2 ms flashes of light (activation of ChR2) on the 16x16 grid as follows: We subdivided the grid into 4x4 blocks. Each such block was treated as one optical stimulus ‘square’. We specified the stimulus ‘density’ as the number of illuminated squares within a block, as a rough correlate of optical intensity. For most runs we used a stimulus density of four. To map to the experiment, we delivered 1, 5 and 15 square stimuli as described in the main section.

To deliver autocorrelated input we reduced the light flash duration to 0.5 ms. We generated autocorrelated spike trains based on the method of (Goldman et al., 1999). In brief we summed multiple instances of a Poisson train, each train shifted by an exponentially decaying distribution of offsets.

All simulations were implemented in MOOSE (Ray & Bhalla, 2008) and the models and figure generation scripts are provided on github/BhallaLab/SynapseVariability. MOOSE uses the Gillespie Stochastic Systems Algorithm (Gillespie, 1977b) for stochastic chemical calculations, and LSODA for deterministic calculations. Large batch simulations were run on a 128 core AMD Epyc 7763 server. We used the Python library *statsmodels* to compute the autocorrelation function.

## Author contributions

Sulu Mohan: Conceptualization, Data curation, Analysis, Investigation, Visualization, Methodology, Writing original draft, Writing review and editing;

Upinder Singh Bhalla: Conceptualization, Resources, Simulation, Visualization, Analysis, Software, Supervision, Funding acquisition, Writing original draft, Project administration, Writing review and editing.

## Acknowledgements

NCBS-TIFR receives the support of the Department of Atomic Energy, Government of India, under Project Identification No. RTI 4006. This project also received support from SERB Core Research Grant, Govt of India, file no. CRG/2022/003135. Animal work in the NCBS/inStem Animal Care and Resource Center was partially supported by the National Mouse Research Resource (NaMoR) grant# BT/PR5981/MED/31/181/2012;2013-2016 & 102/IFD/SAN/5003/2017-2018 from the Indian Department of Biotechnology.

## Supplementary Figures

### Patterned optogenetics reveals diverse STP profiles compared to volley stimulation of SC

Conventionally, electrical stimulation of Schaffer collaterals is done to test CA1 STP. In our experiments, electrical stimulation strength was calibrated to elicit approximately 2-4 mV response in CA1 soma, to keep the stimulus strength comparable between slices (Supplementary Figure 1A,C). We observed short term facilitation in current clamp and short term depression for inhibition in voltage clamp in line with what is reported in the field (Zucker and Regehr 2002, Dobrunz and Stevens 1997, Jackman and Regehr 2017) (Supplementary Figure 1H,I). As electrical stimulation strength gets weaker, the distribution of STP ratios for cells gets broader which is in line with our observations using patterned optical stimulation.

**Supplementary Figure 1:**
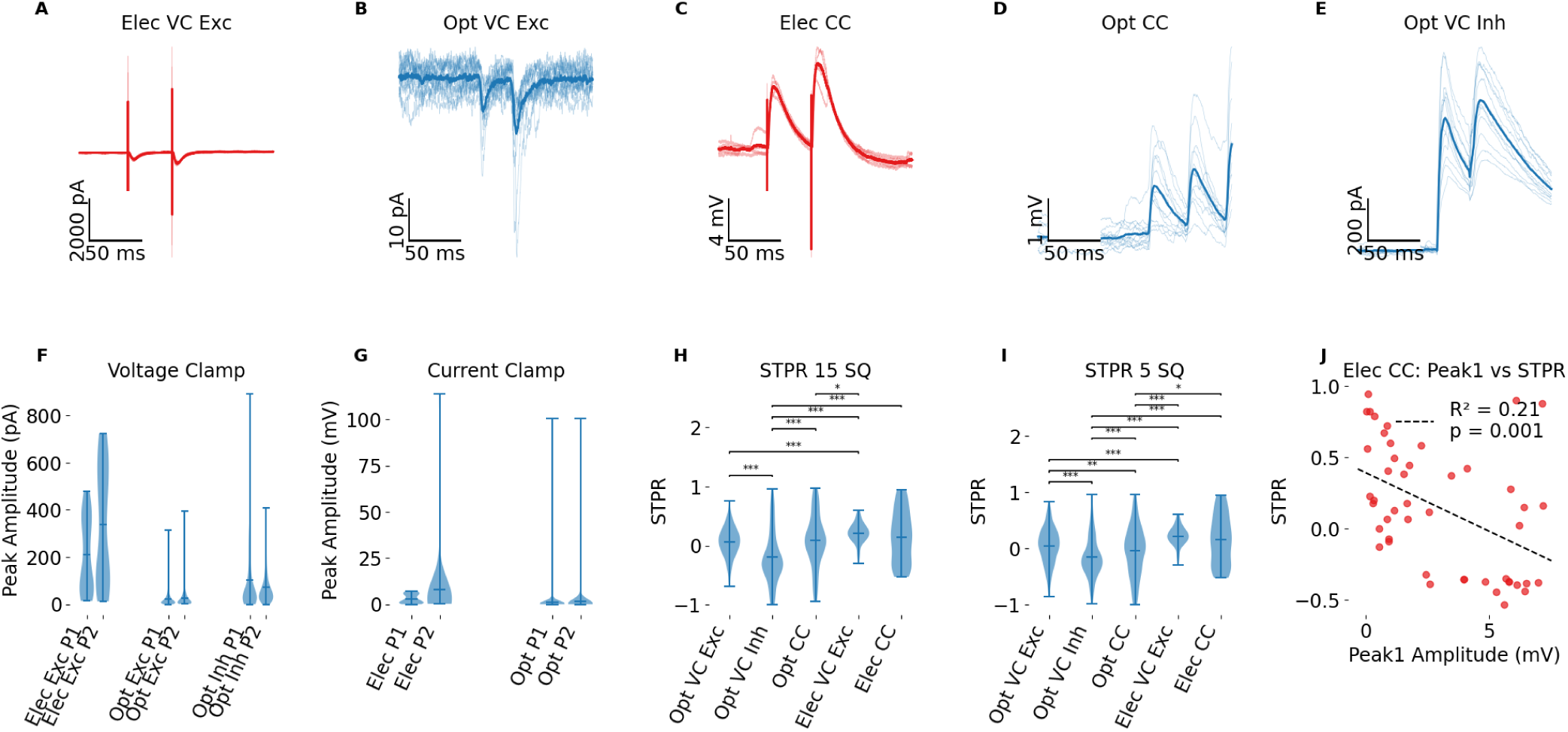
Electrical stimulation of Schaffer collaterals compared with optical stimulation protocols. (A-E) Representative Traces: Example voltage-clamp (VC) and current-clamp (CC) recordings. Bold trace is trial averaged. A. Excitatory postsynaptic currents (EPSCs) evoked by electrical stimulation in voltage-clamp at -70 mV (Elec VC Exc). B. EPSCs evoked by optical stimulation in voltage-clamp at -70 mV (Opt VC Exc). C. Excitatory postsynaptic potentials (EPSPs) evoked by electrical stimulation in current-clamp (Elec CC). D. EPSPs evoked by optical stimulation in current-clamp (Opt CC). E. Inhibitory postsynaptic currents (IPSCs) evoked by optical stimulation in voltage-clamp at 0 mV (Opt VC Inh). (F, G) Violin plots showing the distribution of the first (P1) and second (P2) peak amplitudes for voltage-clamp (F) and current-clamp (G) recordings. (H, I) Violin plots comparing the Short-Term Plasticity Ratio (STPR, calculated as (P2-P1)/(P2+P1)) across all conditions. (H) shows data for 15 SQ patterns, and (I) shows data for 5 SQ patterns. Asterisks denote statistical significance from a Mann-Whitney U test (*p < 0.05, **p < 0.01, ***p < 0.001). (J) Scatter plot illustrating the relationship between the Peak 1 amplitude (mV) and the STPR for the Electrical Current-Clamp (Elec CC) condition. The dashed line represents the linear regression fit, with the R^2^ and p-value shown.

**Supplementary figure 2:**
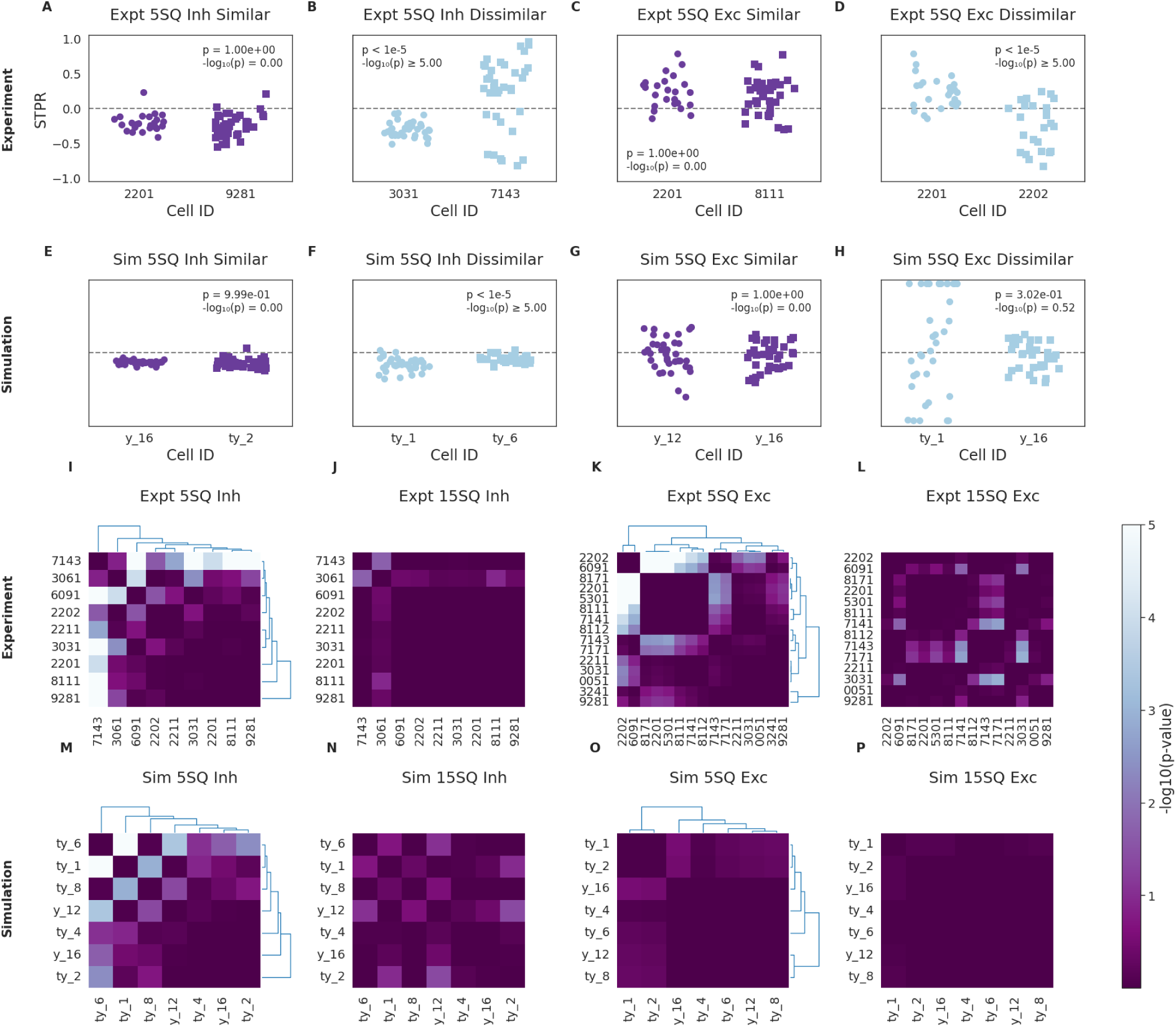
Cell-Pairwise Comparisons of STPR via ANOVA. Pairwise comparisons are performed using one-way ANOVA followed by Tukey’s Honestly Significant Difference (HSD) post-hoc test. The calculated p-values are transformed to -log10(p) and capped at 5.0 (corresponding to p < 1e-5) for visualization. (A-H) Representative Similar and Dissimilar Cell Pairs: The top two rows (Panels A-H) illustrate STPR distributions for representative cell pairs under specific conditions, identified by the ANOVA and Tukey’s HSD analysis as either *Similar* (low -log10(p)) or *Dissimilar* (high -log10(p)). Each stripplot visualizes the distribution of STPR values (Y-axis) for the two selected cells (X-axis). A horizontal dashed grey line at Y=0 indicates STPR=0. Each stripplot includes a text box indicating the p-value from the Tukey HSD test for that specific cell pair, formatted as "-log10(p) = X.XX" or "p < 1e-5 / -log10(p) >= 5.00" if capped. (A-D) Experimental Data: STPR distributions for representative Similar (A) and Dissimilar (B) cell pairs under 5SQ Inhibition. STPR distributions for representative Similar (C) and Dissimilar (D) cell pairs under 5SQ excitation. (E-H) Simulated Data: STPR distributions for representative Similar (E) and Dissimilar (F) cell pairs under 5SQ Inhibition. STPR distributions for representative Similar (G) and Dissimilar (H) cell pairs under 5SQ Excitation. (I-L) Pairwise -log10(p) Heatmaps (ANOVA/Tukey HSD). The bottom two rows (I-P) display heatmaps of the pairwise -log10(p) values (capped at 5.0) between all available cells for various conditions. Brighter colors (darker purple) indicate higher -log10(p) values, signifying greater dissimilarity (stronger statistical difference) between cell pairs. For the 5SQ heatmaps (I, K, M, O), hierarchical clustering (Euclidean distance, Ward linkage) is applied to group cells based on their pairwise p-values. The dendrograms along the top and right of these heatmaps illustrate the clustering structure. For the 15SQ heatmaps (J, L, N, P), the cell order is taken from the corresponding 5SQ heatmap’s clustering to facilitate direct comparison of cell groupings across different numSQ conditions. Color Bar: The common color bar indicates the scale for the -log10(p) values. The scale ranges from vmin_global (the minimum non-zero -log10(p) found across all heatmaps) to vmax_global (the maximum -log10(p) found, or 5.0 if capped). Experimental Data Heatmaps: 5SQ Inhibition (I), 15SQ Inhibition (J), 5SQ Excitation (K), 15SQ Excitation (L) (M-P) Same as I-L for simulated Data: 5SQ Inhibition (M), 15SQ Inhibition (N), 5SQ Excitation (O), 15SQ Excitation (P).

**Supplementary figure 3:**
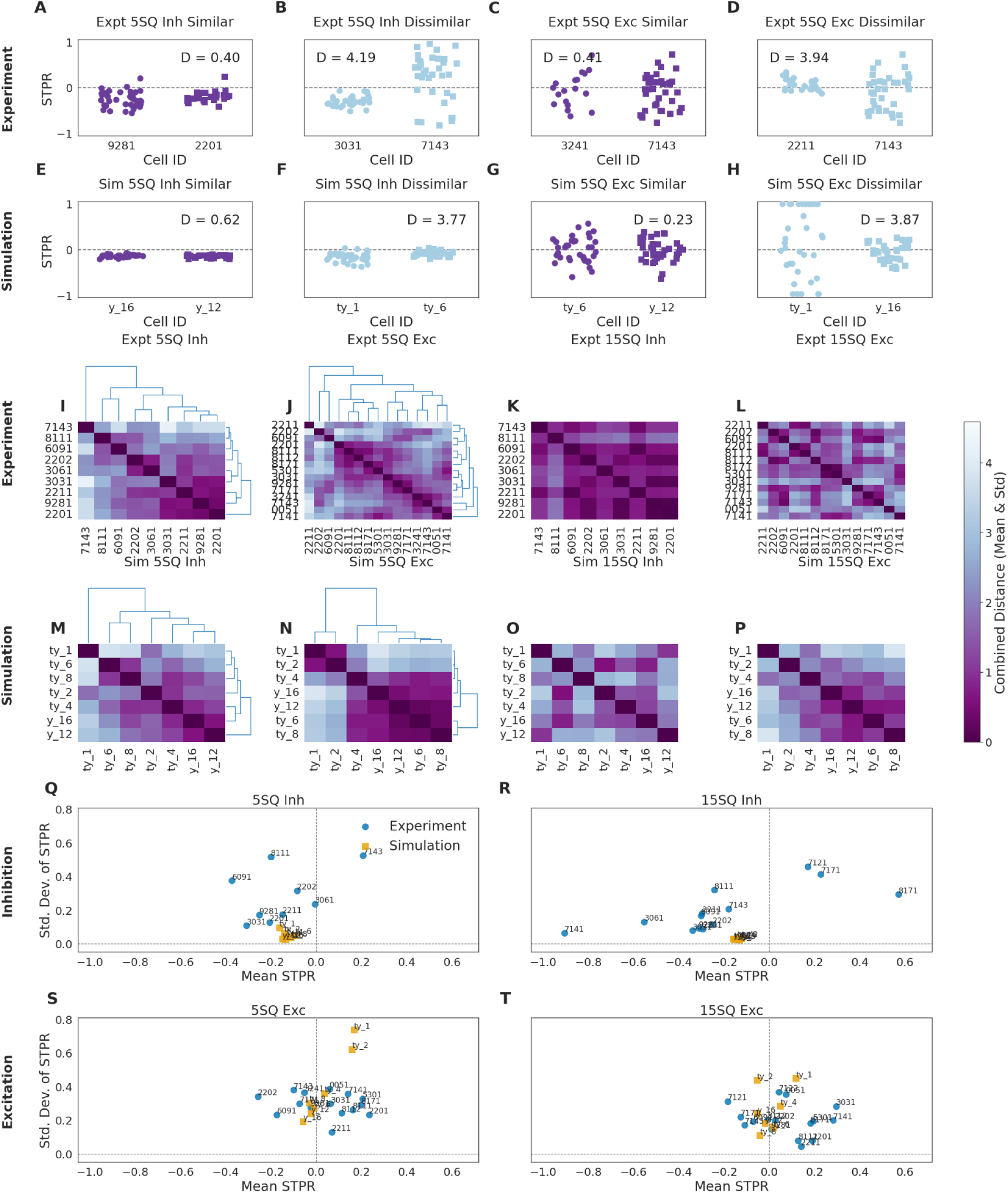
Euclidean distance approach for cell comparisons. (A-H) STPR Distribution and Representative Cell Pairs: Pairwise cell similarity for all cell pair combinations was estimated using a combined distance metric (Euclidean distance on normalized mean and standard deviation of STPR). Panels A–D (Experiment) and E–H (Simulation) show stripplots of STPR values for representative *similar* (left column: A, B, E, F) and *dissimilar* (right column: C, D, G, H) cell pairs under 5-square (5SQ) inhibition (A, C, E, G) and excitation (B, D, F, H). Similarity was assessed using pairwise Euclidean distances (denoted *D*) in a 2D space defined by normalized mean and standard deviation of STPR. (I-P) Cell Similarity Heatmaps and Dendrograms: Panels I-L (Experiment) and M-P (Simulation) show heatmaps of pairwise cell similarity using a combined distance metric (Euclidean distance on normalized mean and standard deviation of STPR). Clustered dendrograms are provided for 5-square conditions (I, J, M, N), with 15-square conditions (K, L, O, P) ordered according to the corresponding 5SQ clustering to highlight consistency. Darker shades indicate greater similarity. Q-T) Mean vs. Standard Deviation Plane: Panels Q-R (Inhibition) and S-T (Excitation) illustrate cell characteristics in the Mean STPR vs. Standard Deviation of STPR plane in both experimental and simulated data. Each point represents a cell, with its short ID labeled. These plots offer a visual representation of the distribution and potential clustering of cell types based on these two key STPR parameters across 5SQ (Q, R) and 15SQ (S, T) inhibition and excitation.

**Supplementary figure 4:**
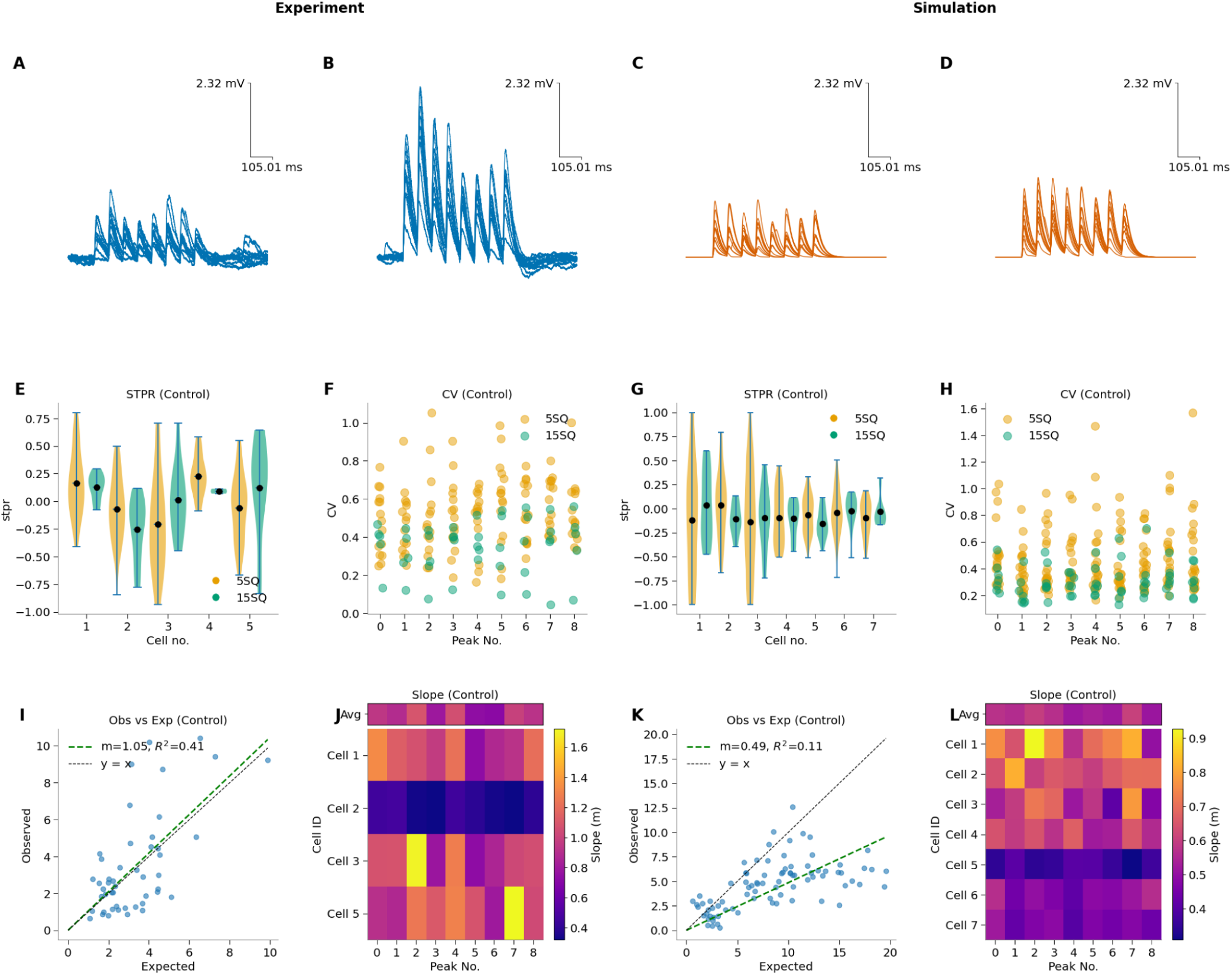
Summation in Current clamp under control and gabazine conditions. (A-D) Representative raw traces from a single cell under current clamp: experimental 5 square (A), experimental 15 square (B), simulation 5 square (C), and simulation 15 square (D). Experimental traces are blue; simulation traces are red. (E, G) Short-Term Plasticity Ratio (STPR) for experimental (E) and simulation (G) data calculated as (peak1-peak2)/(peak1+peak2). Violin plots show the distribution of STPR for 5SQ (orange) and 15SQ (green) inputs. Black dots indicate the mean. (F, H) Coefficient of Variation (CV) of peak amplitudes for experimental (F) and simulation (H) data. Dots represent the CV for each of the first 9 peaks under 5SQ (orange) and 15SQ (green) conditions. (I, K) Synaptic summation linearity for experimental (I) and simulation (K) data. Plots show the observed 15SQ response versus the expected linear sum of three 5SQ responses. The green dashed line is the linear fit to the data; the black dashed line is the unity line (y=x). (J, L) Summation slope heatmaps for experimental (J) and simulation (L) data. Each square shows the summation slope (m) from the linear fit for a specific peak (x-axis) and cell (y-axis). The top ’Avg’ row displays the mean slope for each peak across all cells.

**Supplementary figure 5:**
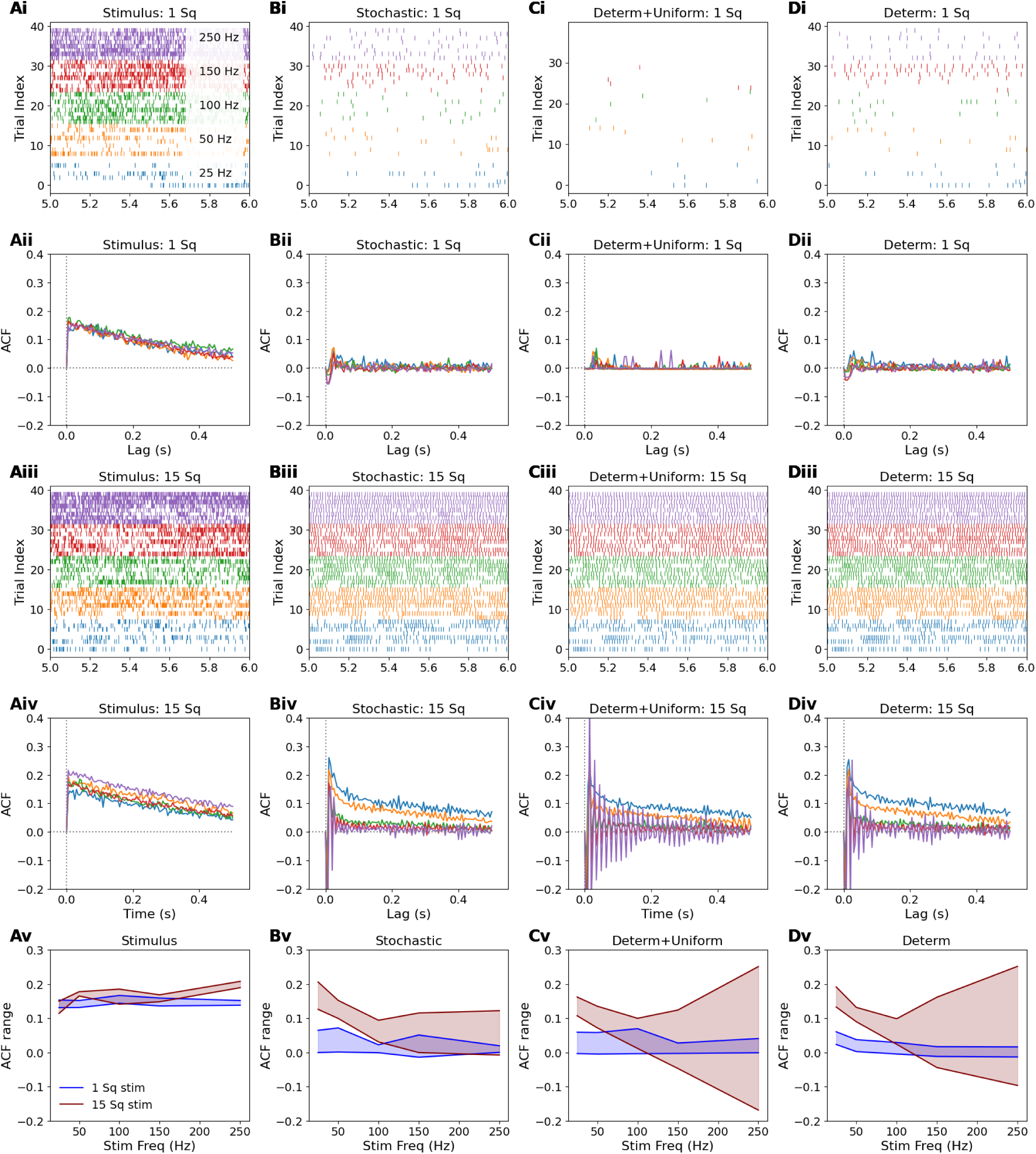
Effect of stochasticity and heterogeneity on output decorrelation. A. Stimulus. Ai: 1-second sample raster of pseudo-random autocorrelated input spike pattern. Different colors indicate different mean frequencies; same colors apply for all plots. Aii: Autocorrelation function (ACF) of the input pattern. Aiii and Aiv: equivalent to Ai and Aii respectively, different seed for 15-square runs. Av: stimulus autocorrelation is nearly independent of frequency. Subsequent runs in B, C and D used the same random number seeds and hence the exact same pseudo-random stimulus spike trains as reported here for A. B. Responses for reference stochastic + heterogeneous model. Bi: Output raster, 1 square stimulus. Bii: Complete decorrelation occurs for 1-square inputs. Biii: Output raster, 15 squares. Biv: Selective decorrelation for inputs above 50 Hz, for 15-square inputs. Bv: There is strong decorrelation except low-frequency 15-square. C. Output spiking patterns and ACFs for deterministic + uniform model. Ci: spiking is low for 1 square. Cii: ACF is flat. Ciii: Spike raster 15 square. Civ: The ACF reports strong periodicity for 150 and 250 Hz inputs which is regular out to almost 200 ms. Cv: Summary plot shows decorrelation for 1 square, but increasing oscillations (positive and negative bounds) at 150 and 250 Hz. D. Responses for model with non-STP synapses, but for deterministic case with heterogeneity retained. Di, Dii: spiking and ACF show complete decorrelation for 1-square input. Diii, Div: frequency-dependent decorrelation, where the cell amplifies the correlation of low-frequency inputs, but reduces it for high-frequencies. Weaker periodicity also emerges for high-frequency inputs. Dv: Decorrelation increases with frequency for 15-square inputs.

Supplementary Data 1:

**Table 1:**
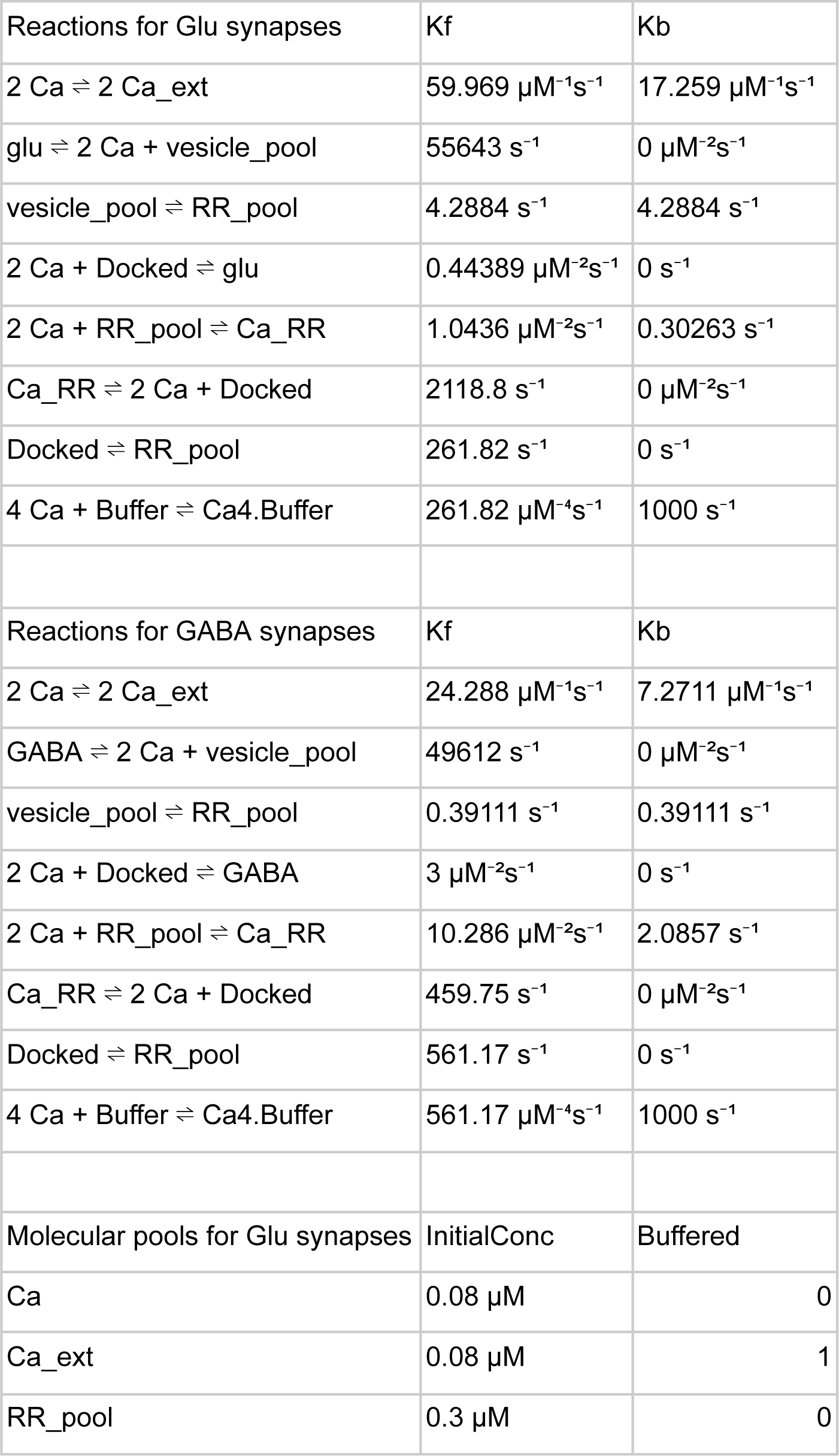
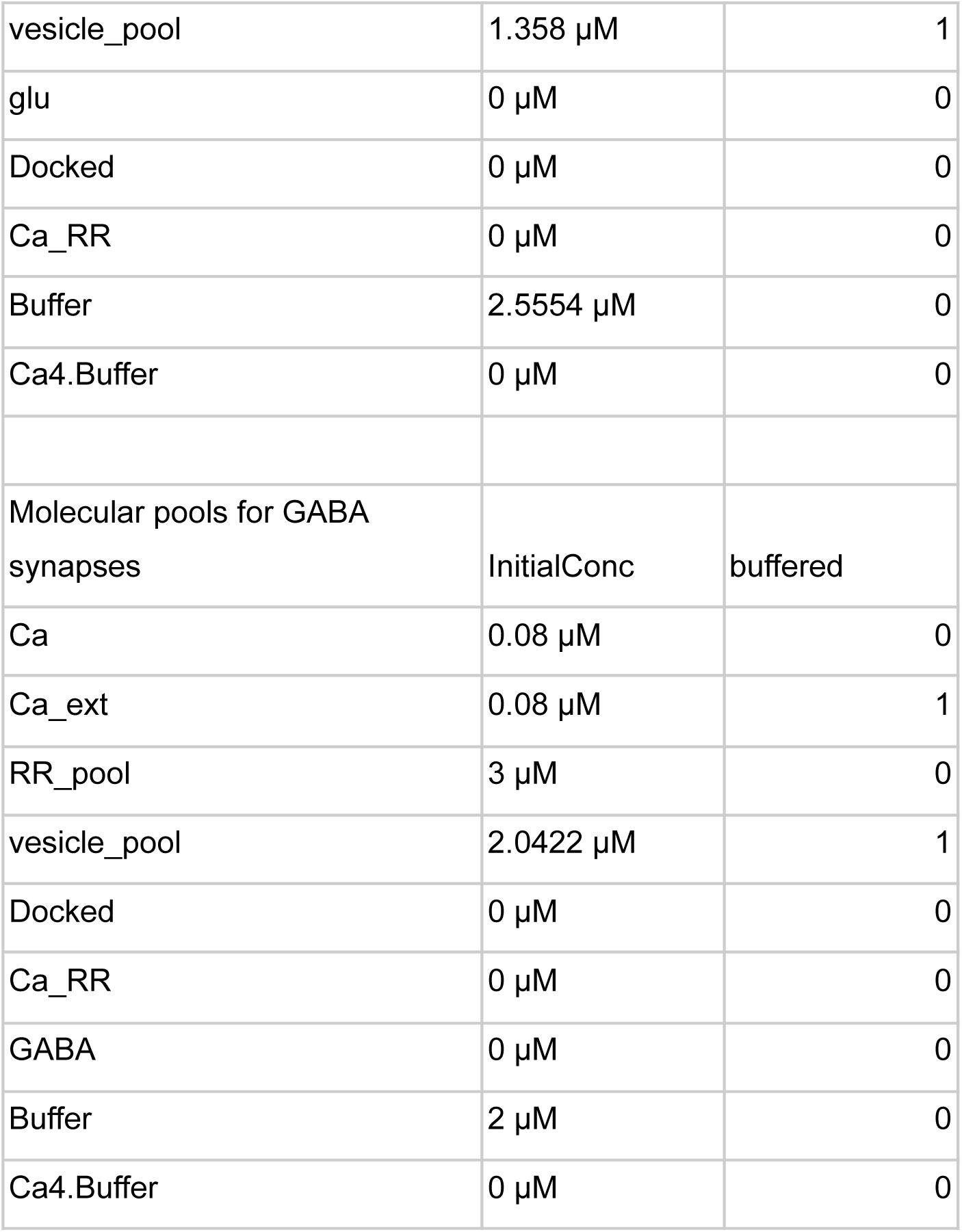
Synapse model parameters.

**Table 2:**
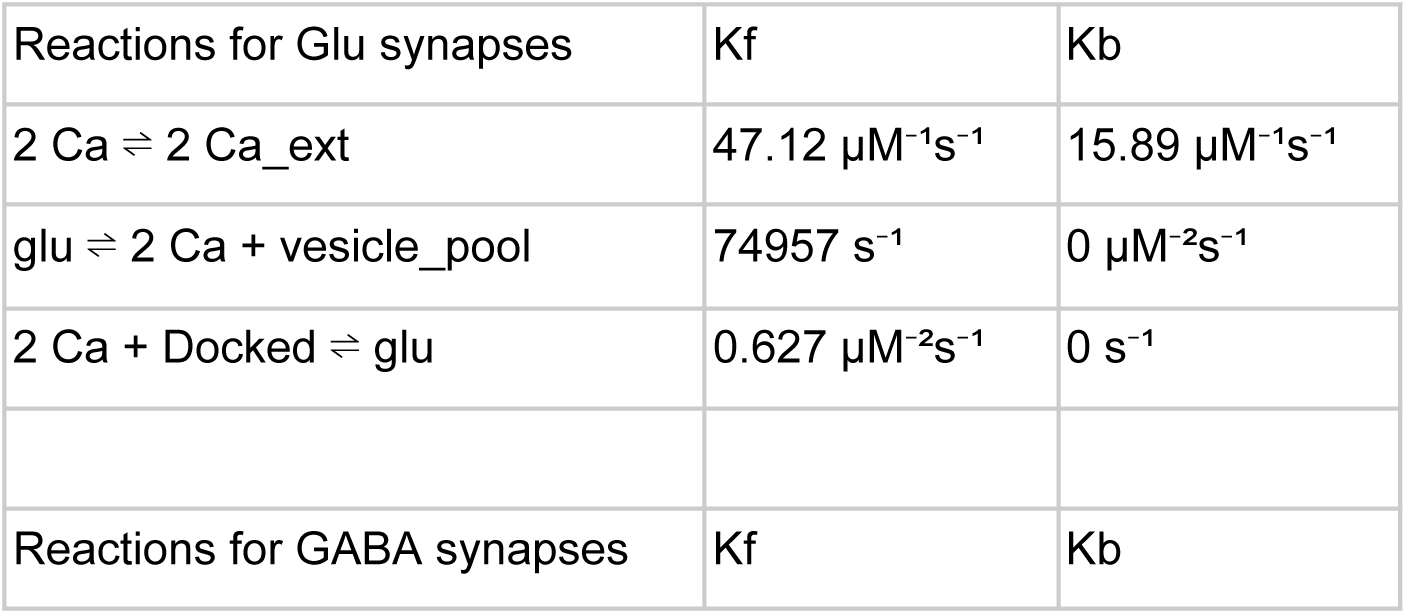
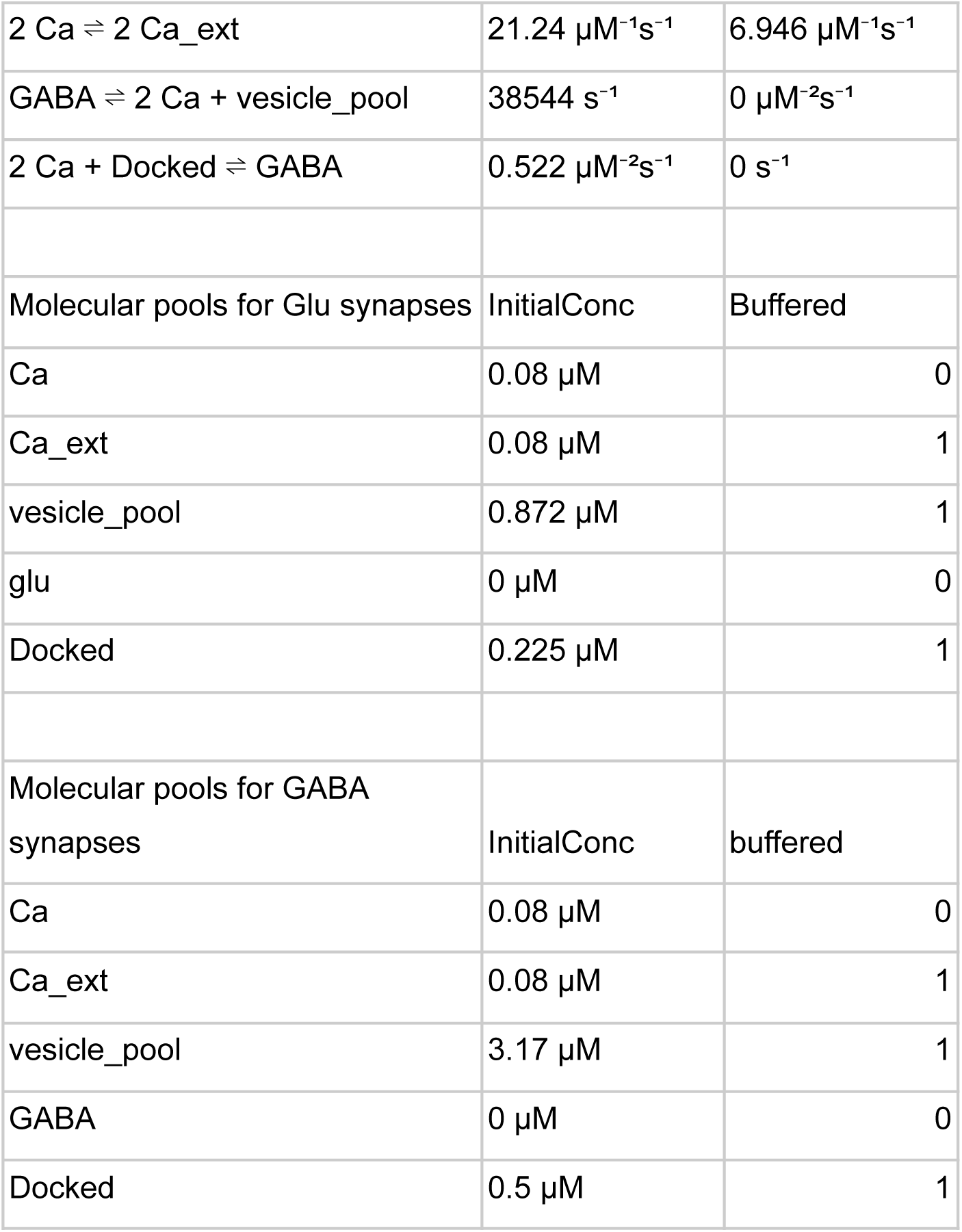
No-STP synapse parameters.

Supplementary Data 2: Electrical Model Parameters

### Electrical Model

The voltage *V* is in mV and referenced to the resting potential. Time is in milliseconds (ms).

#### 1.1 Ion Channel Definitions

These definitions are mostly from (Traub et al., 1991).

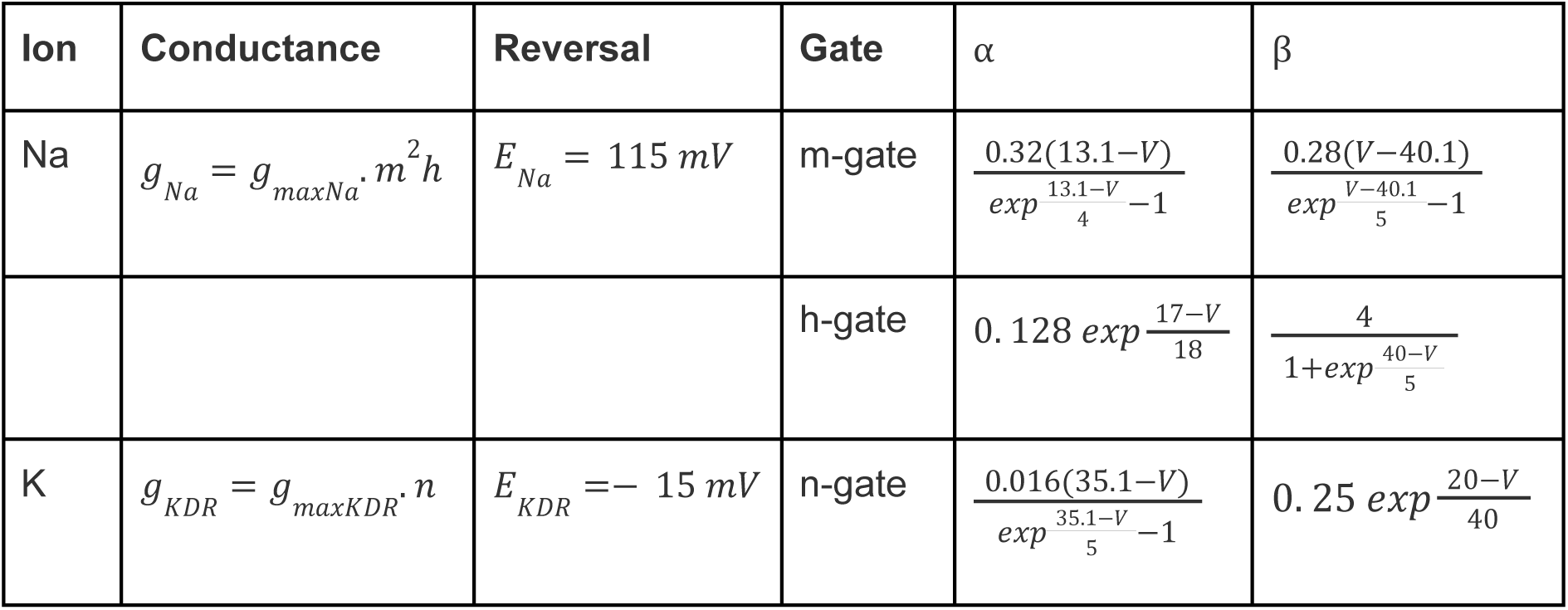

#### 1.2 Receptor-gated Ion Channel Conductances

Glutamate Receptor: AMPA

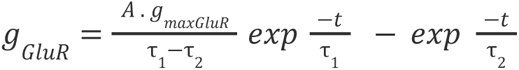

Where: A = normalisation constant such that 𝑔*_Glur_* =𝑔*_maxGlur_* at peak. τ_1_ = 2, τ_2_ = 9

GABA Receptor

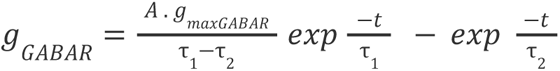

Where: A = normalisation constant such that 𝑔_𝐺𝐴𝐵𝐴𝑅_=𝑔_𝑚𝑎𝑥𝐺𝐴𝐵𝐴𝑅_ at peak. τ_1_= 4, τ_2_ = 9

Glutamate Receptor: NMDA

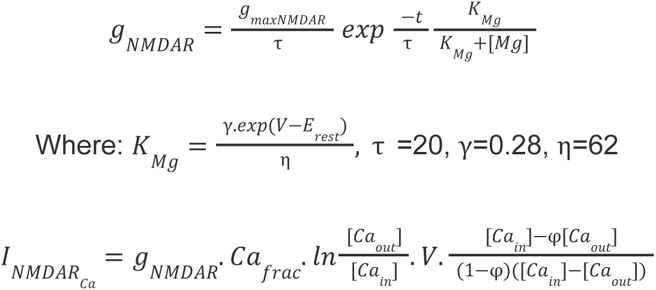

Where: *F* = 96485 sA/mol, *z* = 2, *R* = 8.314 *J/(K.mol)*, *T* = 300 K, φ = exp(*-VFz/RT*), 𝐶𝑎*_frac_* = fraction of current carried at 0 mV by Ca = 0.02, [𝐶𝑎*_out_*] = 1.5 mM, [𝐶𝑎*_in_*] = 0.08 µM

### 1.3 Calcium Pools

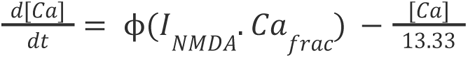

### 1.4 Passive Properties

𝑅_M_ = 1. 0 Ω. 𝑚^2^, 𝑅_A_ = 1. 0 Ω *m^2^, C_M = 0.01 F/m_^2,^ E_rest = −65 mV_*

### 1.5 Channel Distributions

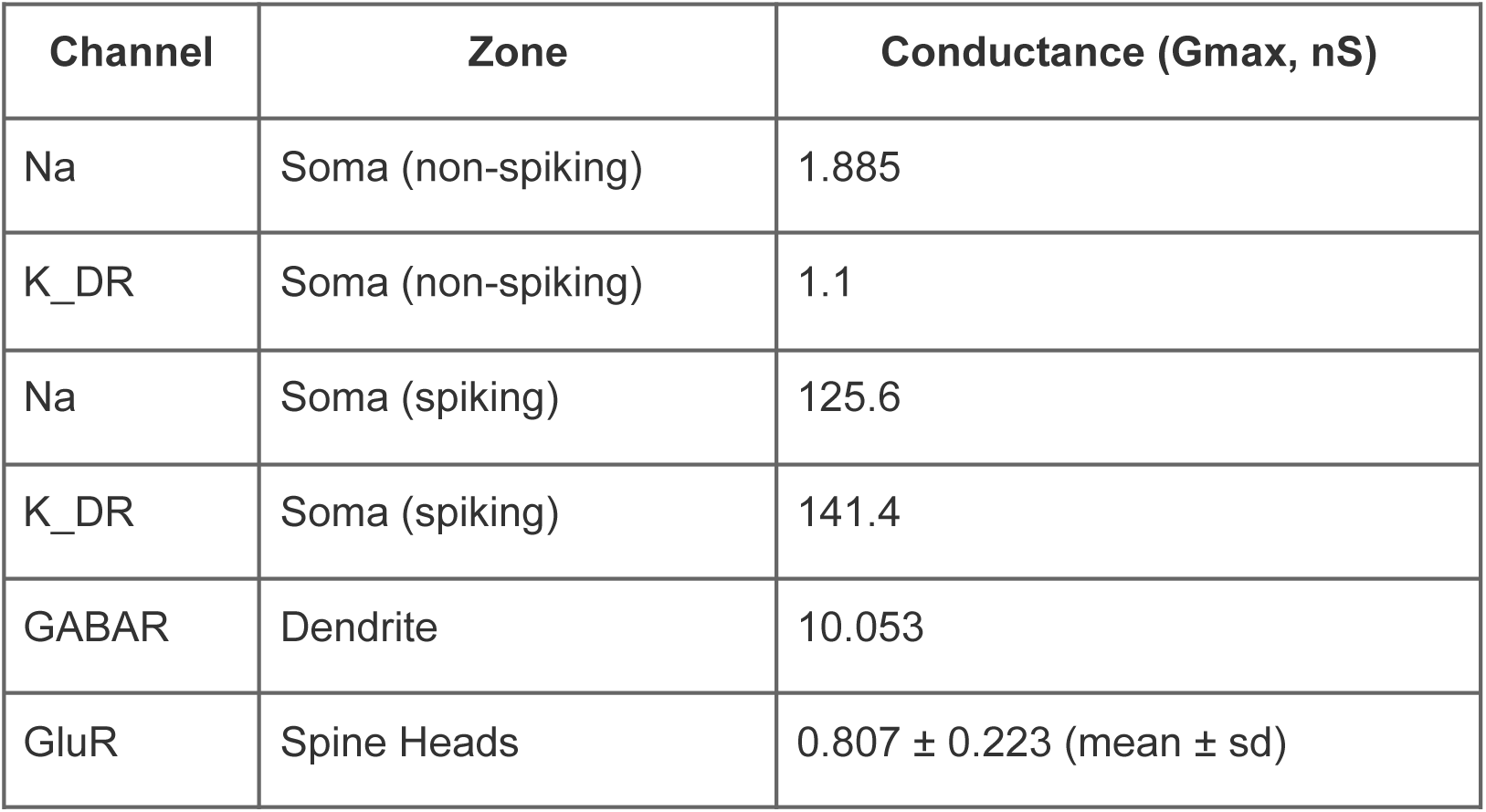
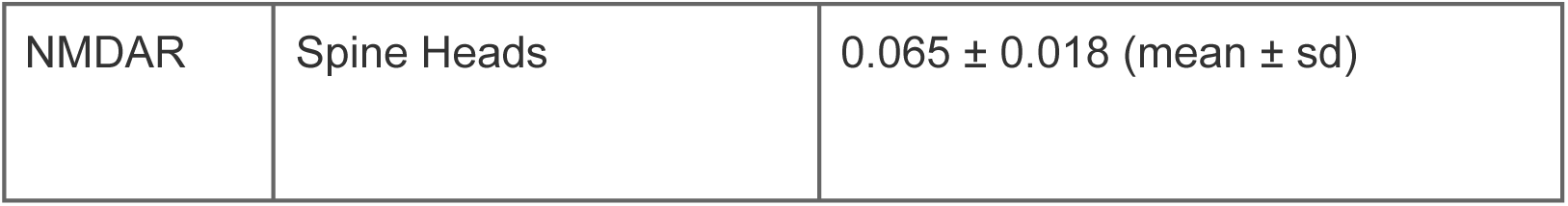

Supplementary Data 3:

Response, variability and heterogeneity aspects of CA1 responses covered in this paper:

**New Insights** , **Expansions**, **Future directions** and **Existing data** beyond the scope of this paper

**Table 1:**
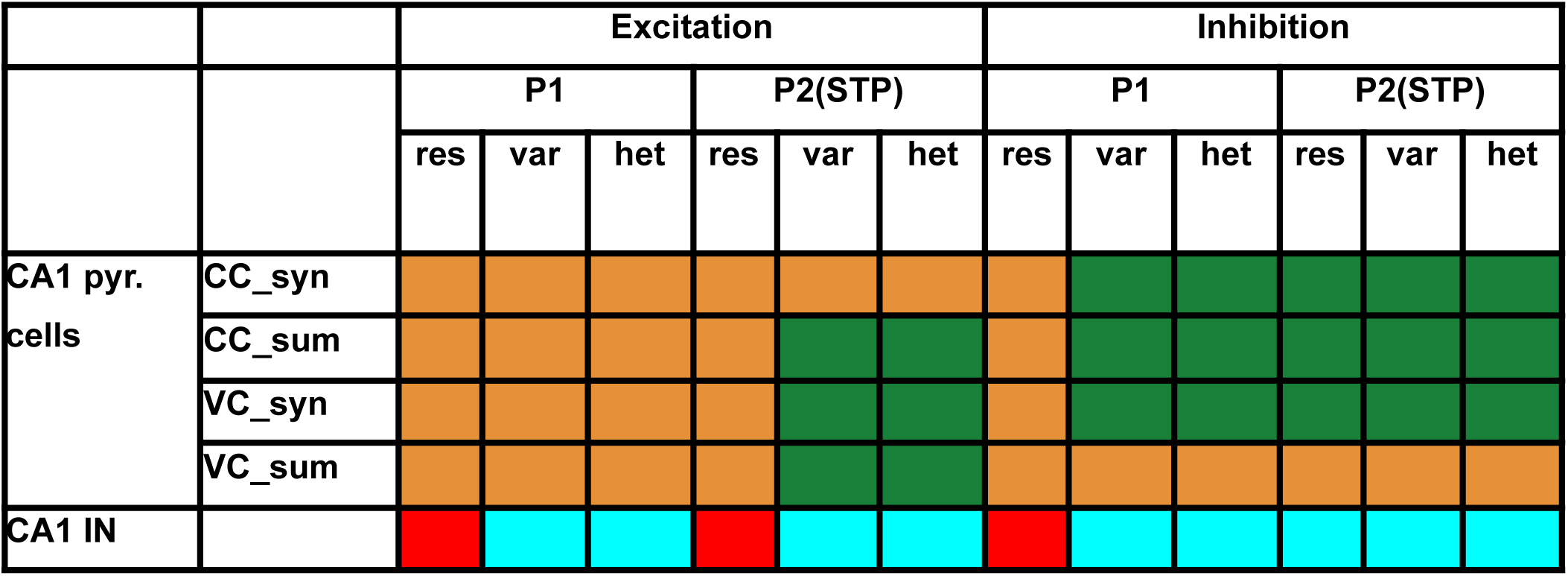
We studied short term plasticity in CA1 pyramidal cells, focusing on response (res), variability (var), and heterogeneity (het) using a paired pulse protocol (P1 and P2). We analyzed both excitation and inhibition at the level of single (syn) and multiple synapses (sum) using current clamp (CC) and voltage clamp (VC) techniques. The table highlights where we have gained new insights (green), expanded on existing literature (yellow), identified gaps in current knowledge (cyan), and acknowledged existing data (red) that are not part of our study. Each combination is color-coded to distinguish these categories.

## Notes

https://github.com/BhallaLab/Cell_Synapse_Variability

### Competing Interest Statement

The authors have declared no competing interest.

