## Supplementary Material for "Heterogeneity and variability in short term synaptic plasticity and implications for signal transformation"

Supplementary Data 1:

Table 1: Synapse model parameters.

| Reactions for Glu synapses | Kf | Kb |
|---|---|---|
| 2 Ca ⇌ 2 Ca_ext | 59.969 $\mu M^{-1}s^{-1}$ | 17.259 $\mu M^{-1}s^{-1}$ |
| glu ⇌ 2 Ca + vesicle_pool | 55643 $s^{-1}$ | 0 $\mu M^{-2}s^{-1}$ |
| vesicle_pool ⇌ RR_pool | 4.2884 $s^{-1}$ | 4.2884 $s^{-1}$ |
| 2 Ca + Docked ⇌ glu | 0.44389 $\mu M^{-2}s^{-1}$ | 0 $s^{-1}$ |
| 2 Ca + RR_pool ⇌ Ca_RR | 1.0436 $\mu M^{-2}s^{-1}$ | 0.30263 $s^{-1}$ |
| Ca_RR ⇌ 2 Ca + Docked | 2118.8 $s^{-1}$ | 0 $\mu M^{-2}s^{-1}$ |
| Docked ⇌ RR_pool | 261.82 $s^{-1}$ | 0 $s^{-1}$ |
| 4 Ca + Buffer ⇌ Ca4.Buffer | 261.82 $\mu M^{-4}s^{-1}$ | 1000 $s^{-1}$ |
| Reactions for GABA synapses | Kf | Kb |
| 2 Ca ⇌ 2 Ca_ext | 24.288 $\mu M^{-1}s^{-1}$ | 7.2711 $\mu M^{-1}s^{-1}$ |
| GABA ⇌ 2 Ca + vesicle_pool | 49612 $s^{-1}$ | 0 $\mu M^{-2}s^{-1}$ |
| vesicle_pool ⇌ RR_pool | 0.39111 $s^{-1}$ | 0.39111 $s^{-1}$ |
| 2 Ca + Docked ⇌ GABA | 3 $\mu M^{-2}s^{-1}$ | 0 $s^{-1}$ |
| 2 Ca + RR_pool ⇌ Ca_RR | 10.286 $\mu M^{-2}s^{-1}$ | 2.0857 $s^{-1}$ |
| Ca_RR ⇌ 2 Ca + Docked | 459.75 $s^{-1}$ | 0 $\mu M^{-2}s^{-1}$ |
| Docked ⇌ RR_pool | 561.17 $s^{-1}$ | 0 $s^{-1}$ |
| 4 Ca + Buffer ⇌ Ca4.Buffer | 561.17 $\mu M^{-4}s^{-1}$ | 1000 $s^{-1}$ |
| Molecular pools for Glu synapses | InitialConc | Buffered |
| Ca | 0.08 μM | 0 |
| Ca_ext | 0.08 μM | 1 |
| RR_pool | 0.3 μM | 0 |
| vesicle_pool | 1.358 μM | 1 |
| glu | 0 μM | 0 |

| | | |
|---|---|---|
| Docked | 0 μM | 0 |
| Ca_RR | 0 μM | 0 |
| Buffer | 2.5554 μM | 0 |
| Ca4.Buffer | 0 μM | 0 |
| Molecular pools for GABA synapses | InitialConc | buffered |
| Ca | 0.08 μM | 0 |
| Ca_ext | 0.08 μM | 1 |
| RR_pool | 3 μM | 0 |
| vesicle_pool | 2.0422 μM | 1 |
| Docked | 0 μM | 0 |
| Ca_RR | 0 μM | 0 |
| GABA | 0 μM | 0 |
| Buffer | 2 μM | 0 |
| Ca4.Buffer | 0 μM | 0 |

Table 2: No-STP synapse parameters.

| Reactions for Glu synapses | Kf | Kb |
|---|---|---|
| 2 Ca ⇌ 2 Ca_ext | 47.12 $\mu M^{-1}s^{-1}$ | 15.89 $\mu M^{-1}s^{-1}$ |
| glu ⇌ 2 Ca + vesicle_pool | 74957 $s^{-1}$ | 0 $\mu M^{-2}s^{-1}$ |
| 2 Ca + Docked ⇌ glu | 0.627 $\mu M^{-2}s^{-1}$ | 0 $s^{-1}$ |
| Reactions for GABA synapses | Kf | Kb |
| 2 Ca ⇌ 2 Ca_ext | 21.24 $\mu M^{-1}s^{-1}$ | 6.946 $\mu M^{-1}s^{-1}$ |
| GABA ⇌ 2 Ca + vesicle_pool | 38544 $s^{-1}$ | 0 $\mu M^{-2}s^{-1}$ |
| 2 Ca + Docked ⇌ GABA | 0.522 $\mu M^{-2}s^{-1}$ | 0 $s^{-1}$ |
| Molecular pools for Glu synapses | InitialConc | Buffered |
| Ca | 0.08 μM | 0 |
| Ca_ext | 0.08 μM | 1 |
| vesicle_pool | 0.872 μM | 1 |

| | | |
|---|---|---|
| glu | 0 µM | 0 |
| Docked | 0.225 µM | 1 |
| Molecular pools for GABA synapses | InitialConc | buffered |
| Ca | 0.08 µM | 0 |
| Ca_ext | 0.08 µM | 1 |
| vesicle_pool | 3.17 µM | 1 |
| GABA | 0 µM | 0 |
| Docked | 0.5 µM | 1 |

Supplementary Data 2: Electrical Model Parameters

**Electrical Model**

The voltage *V* is in mV and referenced to the resting potential. Time is in milliseconds (ms).

### 1.1 Ion Channel Definitions

These definitions are mostly from (Traub et al., 1991).

| **Ion** | **Conductance** | **Reversal** | **Gate** | α | β |
|---|---|---|---|---|---|
| Na | $g_{Na} = g_{maxNa}.m^2h$ | $E_{Na} = 115\ mV$ | m-gate | $\frac{0.32(13.1-V)}{exp\frac{13.1-V}{4}-1}$ | $\frac{0.28(V-40.1)}{exp\frac{V-40.1}{5}-1}$ |
| | | | h-gate | $0.128\ exp\frac{17-V}{18}$ | $\frac{4}{1+exp\frac{40-V}{5}}$ |
| K | $g_{KDR} = g_{maxKDR}.n$ | $E_{KDR} =-\ 15\ mV$ | n-gate | $\frac{0.016(35.1-V)}{exp\frac{35.1-V}{5}-1}$ | $0.25\ exp\frac{20-V}{40}$ |

### 1.2 Receptor-gated Ion Channel Conductances

Glutamate Receptor: AMPA

$$g_{GluR} = \frac{A\ .\ g_{maxGluR}}{\tau_1-\tau_2}\ exp\ \frac{-t}{\tau_1}\ -\ exp\ \frac{-t}{\tau_2}$$

Where: A = normalisation constant such that $g_{GluR}$=$g_{maxGluR}$ at peak. $\tau_1$= 2, $\tau_2$ = 9

GABA Receptor

$$g_{GABAR} = \frac{A\ .\ g_{maxGABAR}}{\tau_1-\tau_2}\ exp\ \frac{-t}{\tau_1}\ -\ exp\ \frac{-t}{\tau_2}$$

Where: A = normalisation constant such that $g_{GABAR}$=$g_{maxGABAR}$ at peak. $\tau_1$= 4, $\tau_2$ = 9

Glutamate Receptor: NMDA

$$g_{NMDAR} = \frac{g_{maxNMDAR}}{\tau}\ exp\ \frac{-t}{\tau}\frac{K_{Mg}}{K_{Mg}+[Mg]}$$

Where: $K_{Mg} = \frac{\gamma.exp(V-E_{rest})}{\eta}$, $\tau$ =20, $\gamma$=0.28, $\eta$=62

$$I_{NMDAR_{Ca}} = g_{NMDAR}.\, Ca_{frac}.\, ln\frac{[Ca_{out}]}{[Ca_{in}]}.\, V.\, \frac{[Ca_{in}]-\varphi[Ca_{out}]}{(1-\varphi)([Ca_{in}]-[Ca_{out}])}$$

Where: *F* = 96485 sA/mol, *z* = 2, *R* = 8.314 *J/(K.mol)*, *T* = 300 K, φ = exp(*-VFz/RT*), $Ca_{frac}$ = fraction of current carried at 0 mV by Ca = 0.02, $[Ca_{out}]$ = 1.5 mM, $[Ca_{in}]$ = 0.08 μM

### 1.3 Calcium Pools

$$\frac{d[Ca]}{dt} = \phi(I_{NMDA}.\, Ca_{frac}) - \frac{[Ca]}{13.33}$$

### 1.4 Passive Properties

$R_M = 1.0\ \Omega.m^2,\ R_A = 1.0\ \Omega.m,\ C_M = 0.01\ F/m^2,\ E_{rest} =- 65\ mV$

### 1.5 Channel Distributions

| Channel | Zone | Conductance (Gmax, nS) |
|---|---|---|
| Na | Soma (non-spiking) | 1.885 |
| K_DR | Soma (non-spiking) | 1.1 |
| Na | Soma (spiking) | 125.6 |
| K_DR | Soma (spiking) | 141.4 |
| GABAR | Dendrite | 10.053 |
| GluR | Spine Heads | 0.807 ± 0.223 (mean ± sd) |
| NMDAR | Spine Heads | 0.065 ± 0.018 (mean ± sd) |
